# Centrosome Loss Alters Axonal and Muscle Development

**DOI:** 10.1101/2025.01.26.634909

**Authors:** Beatriz González, Júlia Sellés-Altés, Jèrica Pla-Parron, Judith Castro-Ribera, Sofia J. Araújo

**Author notes:** Corresponding author. email: S. J. Araújo.

## Abstract

Centrosomes, the primary microtubule-organizing centres (MTOCs), are important during early neuronal development, contributing to polarity establishment and axon formation. Although once considered dispensable in differentiated cells, accumulating evidence suggests that centrosomes remain active in many cell types, with functions that depend on cellular context. In mammalian neurons, centrosome-dependent microtubule (MT) remodelling has been implicated in axon elongation, and centrosome dysfunction has been associated with axonal guidance defects. While active centrosomes have been observed in *Drosophila melanogaster* tracheal cells, their neuronal activity *in vivo* has remained unclear. Here, we examined *Drosophila Sas-4* mutants, which undergo gradual centrosome loss (CL), and found that 50% of homozygous mutants fail to hatch. Analysis revealed the presence of centrosomes in wild-type motor and sensory neurons, with centrosomes localizing near nascent axons in motor neurons. Embryonic CL altered axonal routing and muscle organization, and targeted *Sas-4* reduction in pioneer neurons produced mild guidance phenotypes. Centrosome MTOC activity was confirmed by tubulin localization and MT reassembly analyses. In addition, CL was associated with increased motor axon tortuosity, consistent with a contribution of centrosomes to neuronal morphology. Together, these findings indicate that centrosomes, while not acting as essential MTOCs in this context, contribute to MT dynamics and aspects of early axonal and muscle development.

## Introduction

Centrosomes are small membrane-less organelles that serve as primary microtubule-organizing centres (MTOCs). Composed of a mother-daughter centriole pair surrounded by pericentriolar material (PCM), they anchor and nucleate microtubules (MTs) through y-tubulin ring complexes (y-TuRCs) (Bettencourt-Dias & Glover, 2007). During mitosis, centrosomes organize the spindle apparatus, while their role in interphase is more variable (Conduit *et al*, 2015). Abnormalities in centrosomes are linked to disorders like microcephaly and neurodevelopmental disorders, highlighting the importance of centrosome function in early neurodevelopment (Goundiam & Basto, 2020; Marthiens *et al*, 2013). However, how centrosome numbers affect differentiated cells and their involvement in post-cell-division organogenesis is not yet fully understood. Although traditionally studied in dividing cells, recent insights suggest centrosomes also regulate cellular behaviours in differentiated cells, contradicting the assumption that centrosome-based MTOCs are largely inactive in such contexts.

In vertebrates, centrosomes are critical for embryonic development, often because they are tightly linked to checkpoint controls that ensure accurate cell division and developmental progression (Meyer-Gerards & Bazzi, 2025). Null mutations in mouse *Cenpj* (also called *Sas4*) lead to centriole loss and embryonic arrest at embryonic day 9, and *Sass6*-mutant mouse embryos arrest at mid-gestation (Bazzi & Anderson, 2014; Grzonka & Bazzi, 2024). In contrast, *Drosophila melanogaster* embryos have been reported to develop to adulthood using maternally supplied Sas-4 and Sas-6, despite gradual centrosome loss (CL) (Basto *et al*, 2006; Fatalska *et al*, 2021; Gogendeau & Basto, 2010). However, these adults are sterile, uncoordinated, unable to form cilia, and die shortly after eclosion (Basto *et al*., 2006). Without maternal Sas-4, Sas-4 and Asterless (Asl) *Drosophila* embryonic development arrests early (Stevens *et al*, 2007; Varmark *et al*, 2007), as observed in *C. elegans Sas-4* mutants (Leidel & Gönczy, 2003). In *Drosophila* neural progenitor and epithelial cells, centrosome absence impairs spindle assembly and orientation, leading to defects like neural apoptosis and altered brain size, mimicking either human microcephaly or tumourigenesis (Basto *et al*., 2006; Castellanos *et al*, 2008; Poulton *et al*, 2014; Poulton *et al*, 2017).

Centrosomes are also important for differentiated cellular functions. In mouse fibroblasts, myoblasts and endothelial cells, centrosome position is important for nuclear positioning and cell migration (Chang *et al*, 2015; Gomes *et al*, 2005; Kushner *et al*, 2014). During kidney development, CL leads to decreased branching morphogenesis and small dysplastic kidneys at birth (Cheng *et al*, 2023). In *Drosophila* tracheal terminal cells (TCs), centrosome numbers influence the formation of subcellular lumina during tubulogenesis (Ricolo *et al*, 2016). Increasing centrosome numbers or depleting centrosomal proteins alters this process, stressing centrosome importance in cells with specialized architectures (Ricolo *et al*, 2021). In some neurons, centrosomes play critical roles as MTOCs during early development, particularly in axon formation, a process reliant on microtubule remodelling to establish neuronal polarity (Meka *et al*, 2022). However, other neuronal types need centrosomal microtubule nucleation for migration but not axon formation (Vinopal *et al*, 2023). In mammalian neurons, centrosomes nucleate MTs in the soma, which are subsequently severed and transported into neurites via motor-driven sliding, enabling axon elongation and polarity establishment (Ahmad & Baas, 1995; del Castillo *et al*, 2015; Stiess *et al*, 2010; Yu *et al*, 1993). In hippocampal neurons *in vitro,* suppression of centrosome-mediated functions prevents axonal growth and neurons with more than one centrosome sprout more than one axon (Anda *et al*, 2005). In addition, aberrant centrosome localization, leads to misplaced axonal outgrowth (Anda *et al*, 2010). In human iPSC-derived neurons, depleting centrioles causes significant axonal defects, including mislocalization of axonal MT-associated proteins and affected growth cone morphologies (Lindhout *et al*, 2021). However, as neurons mature, y-tubulin (y-tub) remains essential, but centrosomes lose MTOC activity and become dispensable in mammals and flies (Leask *et al*, 1997; Nguyen *et al*, 2011; Sanchez-Huertas *et al*, 2016; Stiess *et al*., 2010).

These findings collectively suggest that while centrosomes are essential in specific cells, stages and contexts, many differentiated cells can adapt to their loss through alternative MTOCs. However, in many cases, CL induces phenotypes which indicate its important function in cytoskeletal remodelling and the acquisition of specialised cell shapes (Araújo, 2019). To investigate the role of centrosomes in neuronal development, we revisited *Drosophila Sas-4* mutants with reduced centriole numbers and analysed their embryonic development. We quantified the expressivity and penetrance of nervous system phenotypes and examined centriole localization within neurons. Our findings indicate that loss of centrosomal protein function is associated with variable expressivity and penetrance of embryonic phenotypes, which may contribute to inconsistencies among previously reported results. We observed that centrosomes are present and active in *Drosophila* motor and peripheral nervous system (PNS) neurons. Furthermore, centrosome loss (CL) during embryogenesis was associated with axonal misrouting and alterations in muscle development, suggesting a role for centrosomes in the establishment of nervous system and muscle architecture. Consistent with a contribution of centrosomes to neuronal morphogenesis, we identified axonal phenotypes associated with CL.

## Results

### *Sas-4* mutant conditions induce axonal misrouting phenotypes

Spindle assembly abnormal 4 (Sas-4) protein is essential for centriole replication in *Drosophila* and early embryonic development (Basto *et al*., 2006; Stevens *et al*., 2007). However, in contrast to *Sas-4* maternal mutants, *Sas-4* zygotic mutant embryos initially progress through early development due to the maternal contribution, gradually losing centrioles during embryonic and larval stages. This progressive centriole loss makes these mutants an excellent model for studying the role of centrosomes in differentiated cells. By embryonic stages 15–16, centrioles have been reported to be undetectable in 50–80% of cells (Basto *et al*., 2006). By these stages, Sas-4 mutant embryos already exhibit defects in tracheal subcellular lumen formation, indicating that centrioles contribute to the function of some differentiated cells (Ricolo *et al*., 2016). Despite this, Sas-4 homozygous mutants are able to complete development and reach adulthood, although they die soon after eclosion (Basto *et al*., 2006). The observed early adult lethality indicates that loss of Sas-4 affects one or more cellular and developmental processes required for normal viability.

To investigate the role of centrosomes in embryonic development, we first analysed the viability of *Sas-4* loss-of-function (LOF) mutants during embryonic, larval and adult stages. As previously reported, we were able to raise *Sas-4* mutant adults; however, only a proportion of *Sas-4* individuals were found to eclose. These mutants were uncoordinated and became trapped in the food upon eclosion. When isolated at the pupal stage and placed in Petri dishes, they were able to eclose and survive for longer, but without available food, they died within 24-36 hours, which is significantly less than the reported survival rates of unfed flies (2-3 days) (Kopeć, 1928; Pearl & Parker, 1921). These adult individuals exhibited severe motor impairments, barely moving and failing to extend their wings (Fig. 1A). To obtain survival rates, we quantified the percentage of *Sas-4* mutants that reached adulthood, compared to the total number expected, based on Mendelian proportions, from a cross between heterozygous *Sas-4*/*TM3twiGFP* individuals and a cross between *Sas-4*/*TM3twiGFP* and a balanced deficiency, which uncovers the *Sas-4* locus (*Df(3R)BSC221/TM3twiGFP*). Since the homozygous balancer chromosome embryos are early embryonic lethal, we could detect a proportion of 33.3% mutant embryos surviving to embryonic stage 16, corresponding to all *Sas-4* homozygous embryos. However, we only detected 16.7% of *Sas-4* homozygous L1 larvae 36h after egg laying (AEL) (Fig. 1B, n=407), corresponding to 50% of all *Sas-4* homozygous embryos. This proportion was maintained to wandering L3, pupal and adult stages, suggesting that once the barrier of embryonic stages is passed, centrosomes are no longer necessary for viability, and individuals can reach adult stages as previously reported (Basto *et al*., 2006). Nevertheless, our results show that this occurs in only 50% of the *Sas-4* homozygous embryos, indicating that CL induces embryonic lethality in the remaining 50% of cases. Although we used a *Sas-4* mutant chromosome from which all other mutations had been removed by recombination, we also assessed the viability of a transheterozygous combination of *Sas-4* and a deficiency that uncovers the *Sas-4* genomic region. Transheterozygous individuals reached adulthood at a rate of 14.6% (n = 337, Fig. 1B), a result comparable to that observed in *Sas-4* homozygotes and confirming this lethality in *Sas-4* null conditions. We then asked if the *Sas-4* embryos exhibited mutant nervous system phenotypes. To do so, we first analysed FasII positive neurons focusing on the trajectories of their motor axons, particularly on the Intersegmental Nerve (ISN) (Fig. 1C). The penetrance of the ISN phenotype was 52.8% (n=53) quantified as at least 2 defects per embryo, despite variability in the expressivity of the phenotype (Fig. 1E and H and Sup. Fig. 1B and K), which agrees with the larval viability observed. The aberrant axonal pathways included the following: synaptic phenotypes, which were evident when the nerve fibres innervated the wrong target or failed to innervate (Fig. 1E and Sup. Fig. 1B, arrow); misrouting phenotypes, which involved defects in nerves that do not follow the wild-type (wt) stereotypical routes (Fig. 1E and Sup. Fig. 1B, asterisk); and crossing phenotypes, which included aberrant crossing between segments (Fig. 1E arrowhead). The same type of anomalous phenotypes was detected in *Sas-4/Df* mutant embryos (Fig. 1F and H), penetrance 56% (n=25), as well as RNAi *Sas-4* downregulation (Fig. 1G and H), penetrance 33.3% (n=15). We downregulated Sas-4 using a *UAS-Sas-4RNAi* expressed in all post-mitotic neurons, by an elavGal4 driver (Sup. Fig. 1 D and K) and specifically in a subset of neurons, such as the anterior corner cell (aCC), posterior corner cell (pCC) and RP2 pioneers, using RN2-Gal4 (Fujioka *et al*, 2003). Interestingly, when Sas-4 was specifically downregulated only in these neurons, we observed an additional phenotype: fasciculation/defasciculation defects (Fig. 1G arrow) ISN misrouting and defasciculation phenotypes (Fig. 1G and Sup. Fig. 1K, asterisks and arrowhead). In addition, we observed shorter nerve extensions (Fig. 1G, dot), also observed in some *Sas-4* and *Sas-4/Df* mutant embryos (Fig. 1F’ and Sup. Fig. 1C). This suggested these motor axon phenotypes were induced cell-autonomously. We confirmed these downregulation results, using a different RNAi line and could observe similar types of phenotypes (Sup. Fig. 1M-O). Ubiquitous expression of *Sas-4* using a *pUbqSas-4* insertion (Peel *et al*, 2007), could partially rescue the mutant phenotypes, particularly by reducing the number of defects (Fig. 1H and Sup. Fig. 1 E and K).

**Fig. 1.**
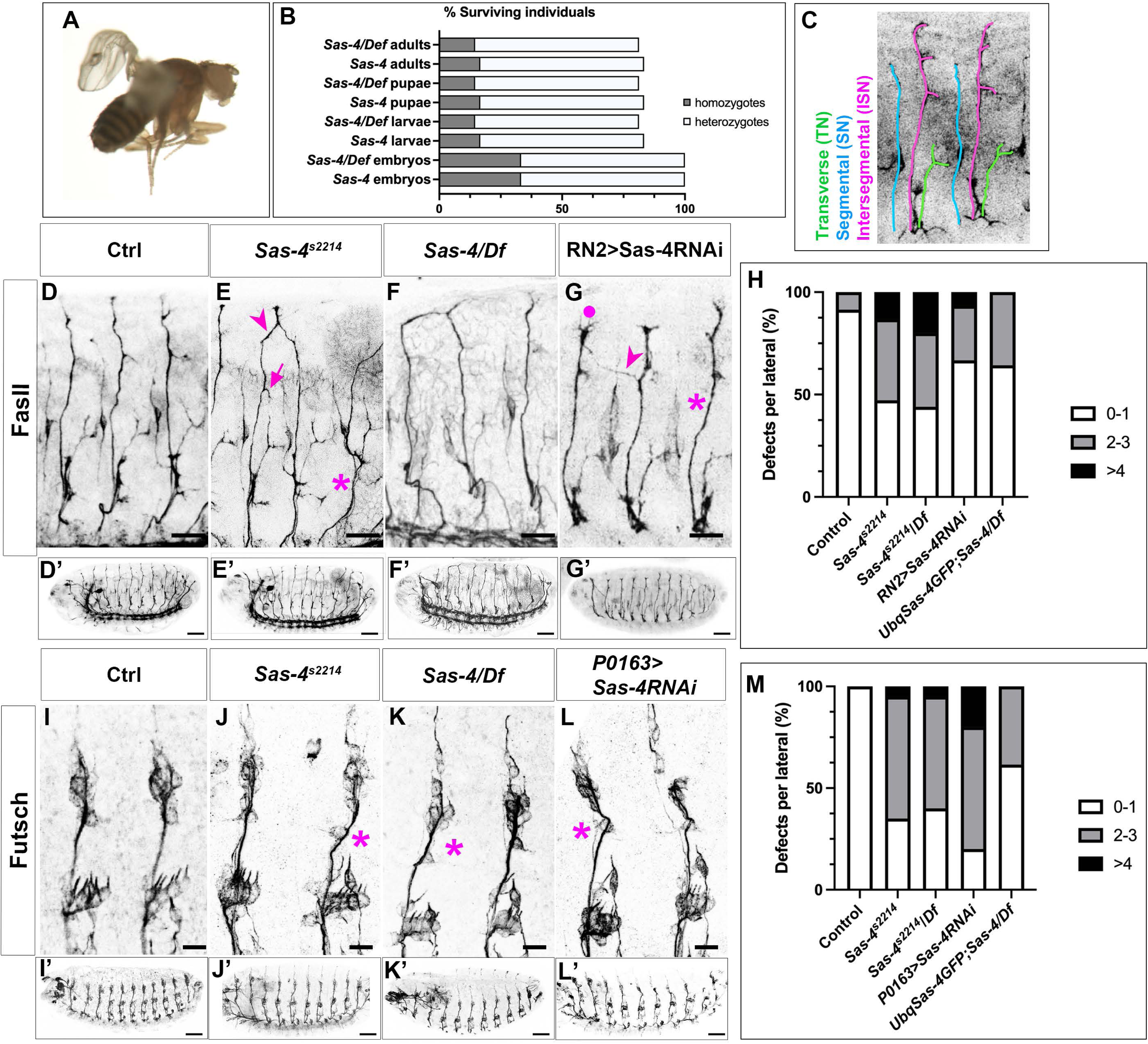
*Sas-4* loss of function induces axonal misrouting defects. (A) *Sas-4* homozygous adult fly at 6h after eclosion. (B) Graphical representation of the percentage of *Sas-4* (n=407) and *Sas-4/Def* (n=337) surviving individuals at different life cycle stages. Homozygous flies that survive to adult stages die after 24-36 hours. (C) Highlight of the three main FasII positive nerves over a microscope image of two embryonic lateral segments. (D-G) Details of three segments from stage 16 embryos stained with anti-FasII antibody; scale bar 10 µm. (D’-G’) Confocal lateral views of stage 16 embryos stained with anti-FasII antibody; scale bar 50 µm. (H) Penetrance and expressivity of the ISN mutant phenotypes in different *Sas-4* mutant conditions; control is *w* (n=35); *Sas-4^s2214^* (n=53); *Sas-4^s2214^*/*Df* (n=25); *RN2Gal4xUASSas-4RNAi* (n=15); *UbqSas-4GFP;Sas-4/Df* (n=28). (I-L) Details of three segments from stage 16 embryos stained with anti-Futsch antibody; scale bar 5 µm (I’-L’) Confocal lateral views of stage 16 embryos stained with anti-Futsch antibody; scale bar 50 µm. (H) Penetrance and expressivity of the sensory axon mutant phenotypes in different *Sas-4* mutant conditions; control is *Sas-4/+* (n=20); *Sas-4^s2214^* (n=20); *Sas-4^s2214^*/*Df* (n=20); *P0163Gal4xUASSas-4RNAi* (n=20); *UbqSas-4GFP;Sas-4/Df* (n=20).

We followed by analysing the peripheral nervous system (PNS) phenotypes of *Sas-4* mutant conditions specifically focusing on their axonal misrouting phenotypes. We could detect axonal misrouting phenotypes in all three mutant conditions: *Sas-4* (penetrance 60% n=20), *Sas-4/Df* (penetrance 55% n=20) and *UAS-Sas-4RNAi* (penetrance 60% n=20) (Fig. 1J-L and M, Sup. Fig. 1F-J) expressed in PNS neurons using *P0163Gal4* (Hummel *et al*, 2000). We could partially rescue these phenotypes by expressing *Sas-4* ubiquitously (Fig.1M and Sup. Fig. 1J and L). Taken together, these results indicate that *Sas-4* mutant and downregulation conditions lead to the generation of mild defects in the nervous system, characterized by incomplete penetrance and variable expressivity. These phenotypes appear to be at least partially cell-autonomous, as Sas-4 downregulation only in motor and sensory neurons results in mild axonal misrouting errors.

### *Sas-4* mutant conditions disrupt muscle development

Since the phenotypes induced by RNAi expression exclusively in neurons did not fully replicate the strength of the motor axon defects observed in *Sas-4* embryos, we next decided to investigate the muscle phenotypes in *Sas-4* mutants. Muscles play a critical role in guiding motor neurons by providing chemical and physical cues essential for accurate axonal pathfinding and target selection (Jeong, 2021). These signals ensure proper neuromuscular connections, essential for coordinated movement. In *Drosophila* mutants lacking muscular structures, such as *twist* (*twi*) embryos, the absence of these cues disrupts motor axon trajectories (Younossi-Hartenstein & Hartenstein, 1993).

In stage 16 *Drosophila* embryos, each abdominal hemisegment (A2–A7) displays a consistent and repeating pattern of 30 somatic muscle fibres, which can be visualized using a myosin-specific antibody (Fig. 2A and C, wt). These 30 syncytial muscle fibres are innervated by two motor nerves: the anterior ISN, which targets the dorsal muscles, and the posterior SN, which innervates the ventral muscles (Fig. 2C, wt). We first analysed the muscle pattern of *Sas-4* mutant embryos and could detect defects that ranged from misplaced to missing muscle fibres (Fig. 2 B, Bߣ). We then analysed these muscle fibre phenotypes in parallel to the axonal defects and were able to correlate many of the axonal defects in *Sas-4* embryos to the location of the muscle pattern disruptions (Fig. 2E’’, asterisk and Sup. Fig. 2B’). In fact, 100% of *Sas-4* embryos with detectable muscle mutant phenotypes also exhibited ISN axonal trajectory disruptions, while 11% of embryos displayed ISN axonal misrouting without any observable muscle phenotypes (n = 27). Accordingly, Sas-4 downregulation in muscle cells, using a *Mef2* driver, induced muscle mutant phenotypes in 80% of embryos, with ISN defects spatially correlating with muscle defects in 63% of the cases (n = 19, Fig. 2G’’ asterisk). Sas-4 downregulation in muscle cells, using a *Mef2* driver together with a *Twist* (*Twi*) driver, induced stronger, more penetrant ISN phenotypes that were dose dependent (Sup. Fig. 2E, F) Consistent with these findings, downregulation of Sas-4 in pioneer neurons using the RN2 driver resulted in axonal misrouting phenotypes in 33% of the embryos (n = 15) and very mild muscle mutant phenotypes in only 20% of the embryos (n = 10). Collectively, these findings suggest that the *Sas-4* axonal misrouting phenotypes arise from two distinct contributions: an autonomous component originating from their own CL and a non-autonomous one driven by CL-affected muscle cells (Fig. 2C, *Sas-4*).

**Fig. 2.**
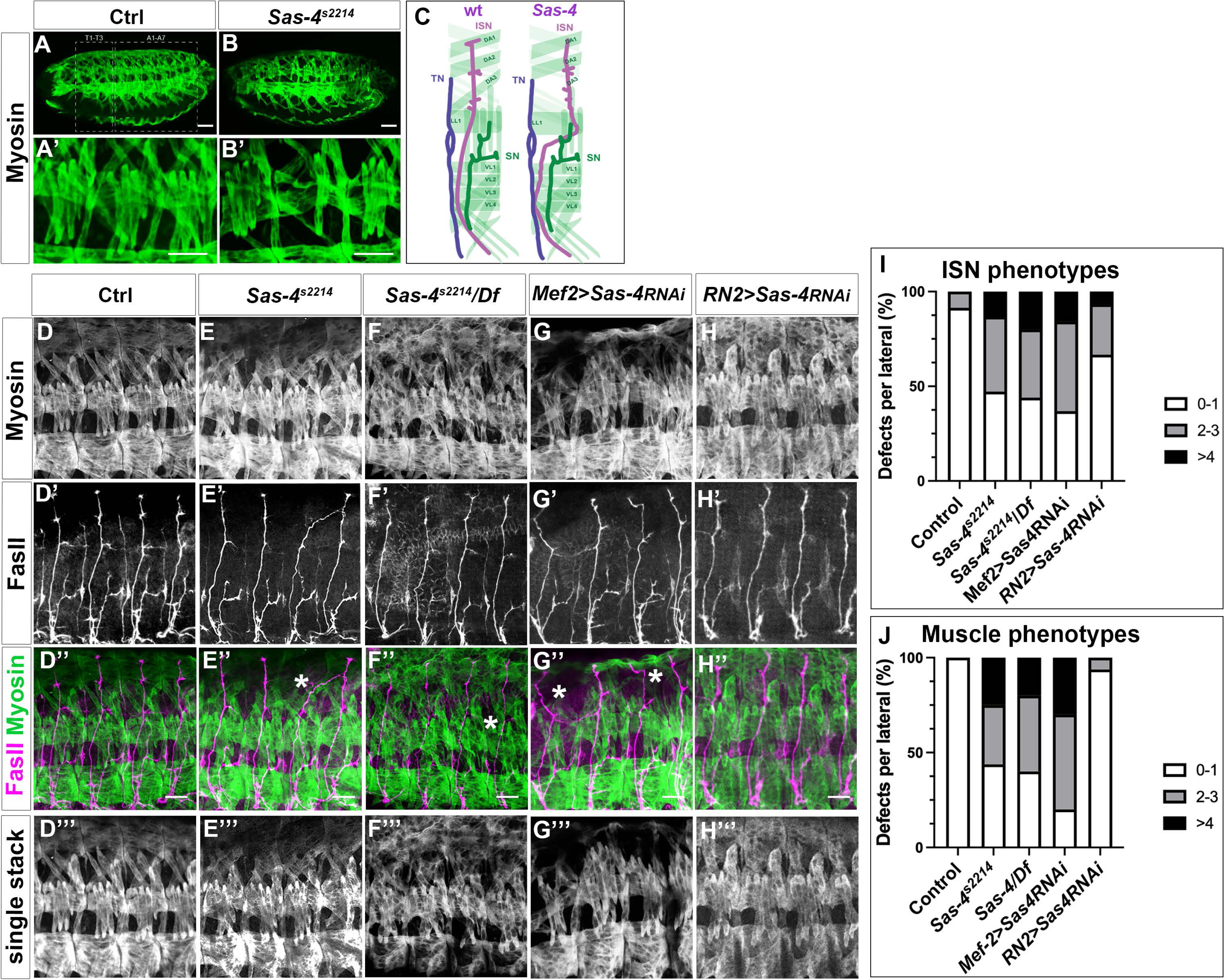
*Sas-4* loss of function induces muscle defects. (A,B) Stage 16 embryos stained with an antibody against myosin to mark the embryonic muscle structure in green. (A-A’) Control embryo and (B-B’) *Sas-4* mutant embryo. (C) Schematic representation of the motor axon muscle innervation pattern of a control (*w*) and *Sas-4* mutant embryo. TN, transverse nerve; ISN, intersegmental nerve, and SN, segmental nerve. (D-H) Confocal projections of four segments from stage 16 embryos of different mutant conditions stained with anti-myosin (green and D’) and anti-FasII (magenta and D’’) antibodies; scale bar 10 µm. (D’’-H’’) composite images with the two channels (anti-myosin in green and anti-FasII in magenta). (D’’’-H’’’) Myosin-stained muscle single-stacks. Scale bar 10 µm. (I) Penetrance and expressivity of the ISN mutant phenotypes in different *Sas-4* mutant conditions; control is *Sas-4/+* (n=27); *Sas-4^s2214^* (n=27); *Sas-4^s2214^*/*Df* (n=25); *Mef2Gal4xUASSas-4RNAi* (n=19); *RN2Gal4xUASSas-4RNAi* (n=15). (J) Penetrance and expressivity of the muscle mutant phenotypes in different *Sas-4* mutant conditions; control is *w* (n=16); *Sas-4^s2214^* (n=16); *Sas-4^s2214^*/*Df* (n=10); *Mef2Gal4xUASSas-4RNAi* (n=10); *RN2Gal4xUASSas-4RNAi* (n=16).

### *Sas-6* mutants have axonal guidance and muscle developmental defects

Centrosome assembly depends on a small set of conserved proteins, the absence of any one of which results in CL; *Sas-4* and *Sas-6* are key components of this process (Gartenmann *et al*, 2020). To rule out the possibility that the observed axonal and muscle defects were specifically due to mutations in *Sas-4* and not CL, we continued by analysing the viability of *Sas-6* LOF mutants during the embryonic, larval, and adult stages. We used a CRISPR-Cas9 generated *Sas-6* induced deletion mutant, *Sas-6^Δb^* (Z. Novak and J. Raff, unpublished). We successfully raised *Sas-6* adults (Fig. 3A), which, like *Sas-4* adults, died within 24-36 hours (Fig. 1A). To corroborate that this *Sas-6* allele induced CL in embryos, we counted the number of centrosomes in *Sas-6^Δb^* embryos and observed a reduction at stage 16 (Fig. 3B). Like *Sas-4* mutants, 33.3% of *Sas-6^Δb^* and *Sas-6^Δb^/Def* embryos reached embryonic stage 16, corresponding to all *Sas-6^Δb^* homozygous and *Sas-6^Δb^/Def* embryos. We only detected 11.7% (n=571) of *Sas-6^Δb^* and 16.8% (n=486) of *Sas-6^Δb^/Def* homozygous L1 larvae 36h after egg laying (AEL) (Fig. 3C), corresponding to 35% of all *Sas-6* homozygous individuals and 50% of *Sas-6^Δb^/Def* individuals. This proportion was maintained through the wandering L3, pupal, and adult stages, suggesting that, like in *Sas-4* mutants, centrosomes seem not to be essential for larval and pupal development once the embryonic stages are completed.

**Fig. 3.**
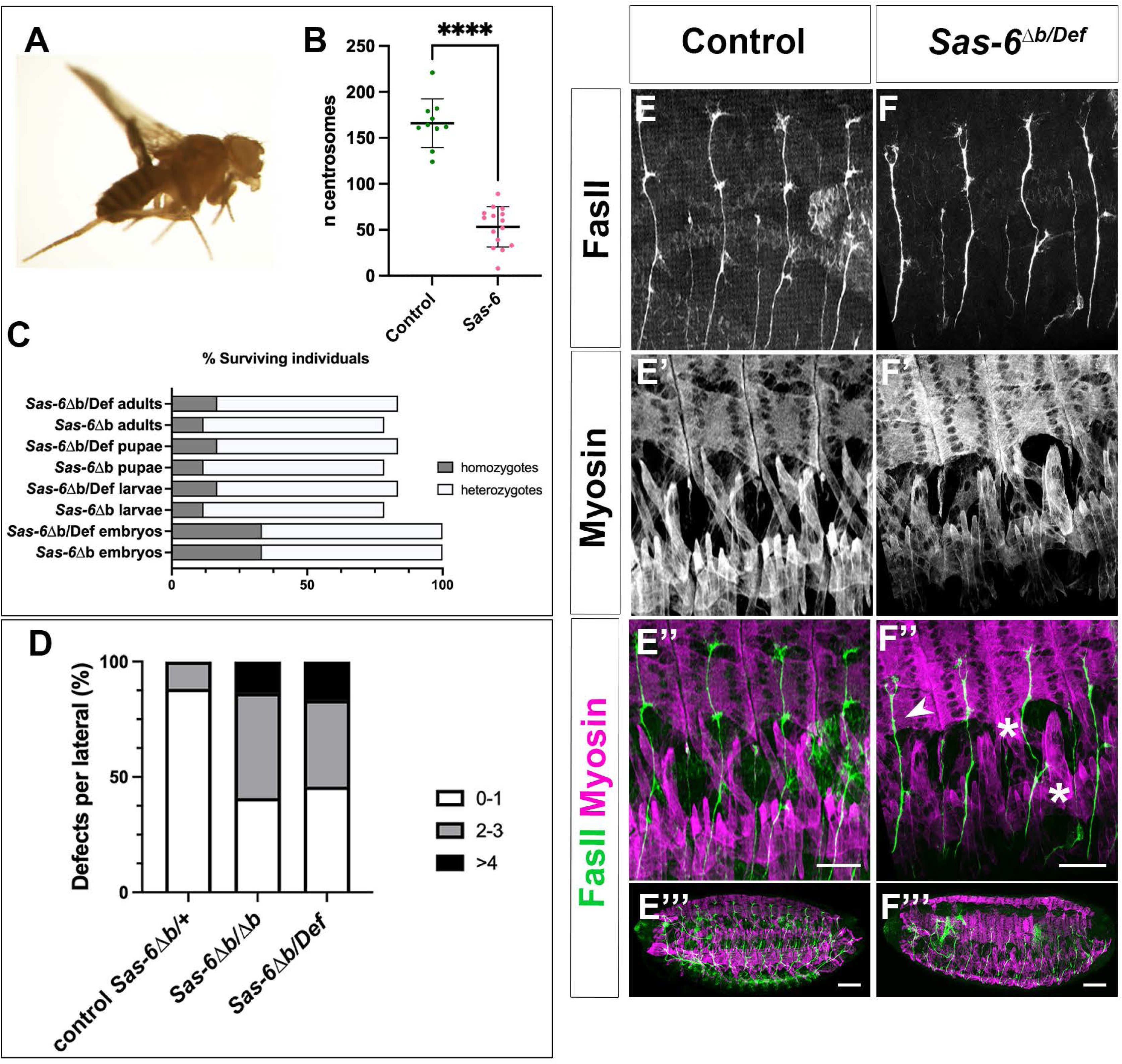
*Sas-6* loss of function induces axonal misrouting and muscle defects. (A) *Sas-6* homozygous adult fly at 6h after eclosion. (B) Quantification of the number of centrosomes per stage 16 embryonic segmental area in Control, *Sas-6^Δb^/+* embryos, (mean 165.9 ± 26.4, n=10) and *Sas-6^Δb^* embryos displaying PNS mutant phenotypes (mean 53.2 ± 21.9, n=15); unpaired t-test with Welch’s correction p<0.0001. (C) Graphical representation of the percentage of *Sas-6^Δb^*/*Sas-6^Δb^* (n=571) and *Sas-6^Δb^*/Def (n=486) surviving individuals at different life-cycle stages. (D) Penetrance and expressivity of the ISN mutant embryonic phenotypes in *Sas-6^Δb^* and *Sas-6^Δb^*/Def embryos; controls are heterozygous *Sas-6^Δb^/+* (n=17); mutants are *Sas-6^Δb/Δb^* (n=22) and *Sas-6^Δb^*/Def (n=24). (E, F) Confocal projections of four segments from stage 16 embryos of control and *Sas-6^Δb^*/Def mutants stained with anti-myosin (magenta and E’, F’) and anti-FasII (green and E, F) antibodies. (E’’-F’’) Composite projections with the two channels; scale bar is 10 µm. (E’’’-F’’’) Composite projections of the two channels showing the whole embryo; scale bar is 50µm.

We then investigated whether *Sas-6* embryos exhibited mutant axonal guidance and muscle phenotypes. We observed ISN axonal misrouting in *Sas-6^Δb^* and *Sas-6^Δb^/Def* embryos (Fig. 3D, F), with a penetrance of 59.1% (n=22) for *Sas-6^Δb^* and 54.1% (n=24) for *Sas-6^Δb^/Def*, similar to what we observed in *Sas-4* mutant conditions. The severity of the defects was also like *Sas-4* mutant conditions, with axonal misrouting (Fig. 3F’’, asterisks) and synapse (Fig. F’’, arrowhead) phenotypes. In most cases, the ISN axonal misrouting also correlated with muscle phenotypes (Fig. 3E’’ and F’’, asterisks). Taken together, these findings suggest that CL through the loss of *Sas-4* or *Sas-6* can affect motor neuron axon guidance.

### Pioneer neurons have centrosomes localizing to the sites of axon outgrowth

In *Drosophila*, the aCC and RP2 motor neurons are pioneer neurons which influence growth of the ISN (Sánchez-Soriano & Prokop, 2005) (Fig. 4 A). Pioneer neurons extend growth cones that navigate a future fibre tract and must interpret the full complexity of environmental cues to define the path of the nerve (Harrison, 1910; Jacobs & Goodman, 1989). Therefore, we investigated whether centrosomes could be detected in the soma of these neurons in stage 12-13 when they are starting to extend their axons. We used an antibody against Pericentrin-like protein (Plp, CP309) to detect the centrosome and we showed that this colocalized with Asl in most cases (Sup. Fig. 3, 89.7% of neuronal Asl foci colocalize with Plp, and 84.8% of neuronal Plp foci colocalize with Asl; n=60 neurons) making it a good tool to detect centrosomes in these cells. We found that centrosomes were present in the soma of both aCC (Fig. 4 B and C) and RP2 motor neurons (Fig. 4 D). Specifically, in the case of the aCC, a centrosome was observed in the quadrant where the axon exit point is first detected in 72% of the cases (Fig. 4B n=75). Additionally, we did not detect centrosomes in the quadrant opposite the axon exit point (Fig. 4B n=75). In *Sas-4* mutants, centrosomes were largely undetected in both aCC/pCC neurons and RP2 neurons. (Fig. 4 E, F). At these stages, the aCC and pCC neurons are very close together, making it difficult to differentiate between the somas of the two neurons. Therefore, we quantified the number of centrosomes in aCC+pCC and RP2 neurons in both control and *Sas-4* mutant embryos. In control embryos, we observed around 2.53 centrosomes per pair of aCC and pCC neurons, while in *Sas-4* mutant, this number was significantly reduced to 0.81 (Fig. 4G n=76). Similarly, for RP2 neurons, control embryos exhibited 0.81 centrosomes per RP2 neuron, whereas *Sas-4* embryos showed a reduction to 0.26 per neuron (Fig. 4G n=76). The observed decrease in centrosome number is in line with the proposed role of Sas-4 in centriole formation in pioneer neurons of the ventral nerve cord (VNC) during *Drosophila* embryogenesis.

**Fig. 4.**
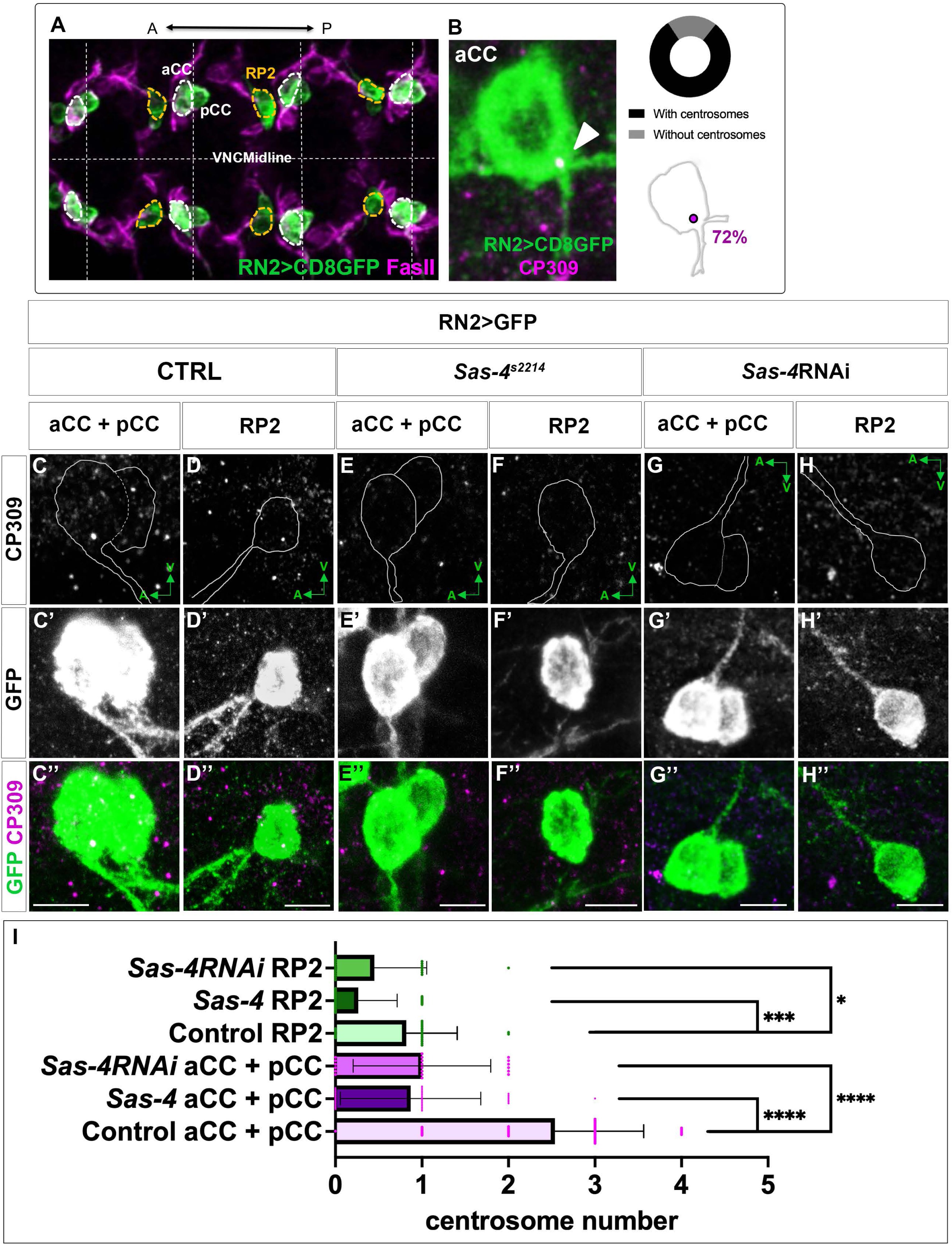
Pioneer neurons have centrosomes localizing to the sites of axon outgrowth. (A) Confocal image showing 3 segments of the embryonic Ventral Nerve Cord (VNC) midline marked with an antibody against FasII and GFP driven by the RN2 driver marking the aCC, pCC and RP2 in green. (B) Confocal section showing an aCC neuron and its centriole at the axonal exit point, marked with an antibody against Plp (CP309); quantification of centrioles present in different aCC quadrants and the percentage of aCC with centrioles present at the axonal exit point (72%, n=75). (C, D) Plp (CP309) marking all centrosomes and the neuronal body drawn from the channel marking the neuron (C’, D’). (C’’) aCC+pCC and RP2 (D’’) marked in green and Plp (CP309) in magenta. Scale bar is 5µm. (G) Quantification of centriole numbers in control and *Sas-4* aCC+pCC and RP2 pioneer neurons; Control RP2 average 0.81 (n=22), control aCC+pCC average 2.53 (n=41), *Sas-4* RP2 average 0.26 (n=34) and *Sas-4* aCC+pCC average 0.87 (n=38); unpaired t-test with Welch’s correction control RP2 vs *Sas-4* RP2 p=0.0005, control aCC+pCC vs *Sas-4* aCC+pCC p<0.0001.

### Pioneer motor neurons have active centrosomes

After revealing that ISN pioneers maintain their centrosomes, we asked whether these could be active MTOCs. To determine if the centriole could organize a functional centrosome and serve as a major site of neuronal microtubule nucleation in aCC and RP2 neurons, we analysed the localization of acetylated-tubulin (ace-tub) and y-tubulin (y-tub). MTs are nucleated and anchored by the microtubule nucleator y-tub, which is embedded within the pericentriolar material (PCM) of the centrosome (Lin *et al*, 2015). In 69.4% of aCC/pCC and 68.8% of RP2 neurons we could observe an overlap between stable acetylated MTs and Plp positive centrosomes (Fig. 5A, B arrowheads and E). In addition, in 57.5% of aCC/pCC and 46.1% of RP2 neurons we could observe an overlap between y-tubulin and Plp positive centrosomes (Fig. 5C, D, arrowheads and E). This suggests that there are centrioles in these neurons which are not active as MTOCs at these stages (Fig. 5C’’ asterisk). In conclusion, these results indicate that at least some MT organization in pioneer neurons is dependent on the centrosome.

**Fig. 5.**
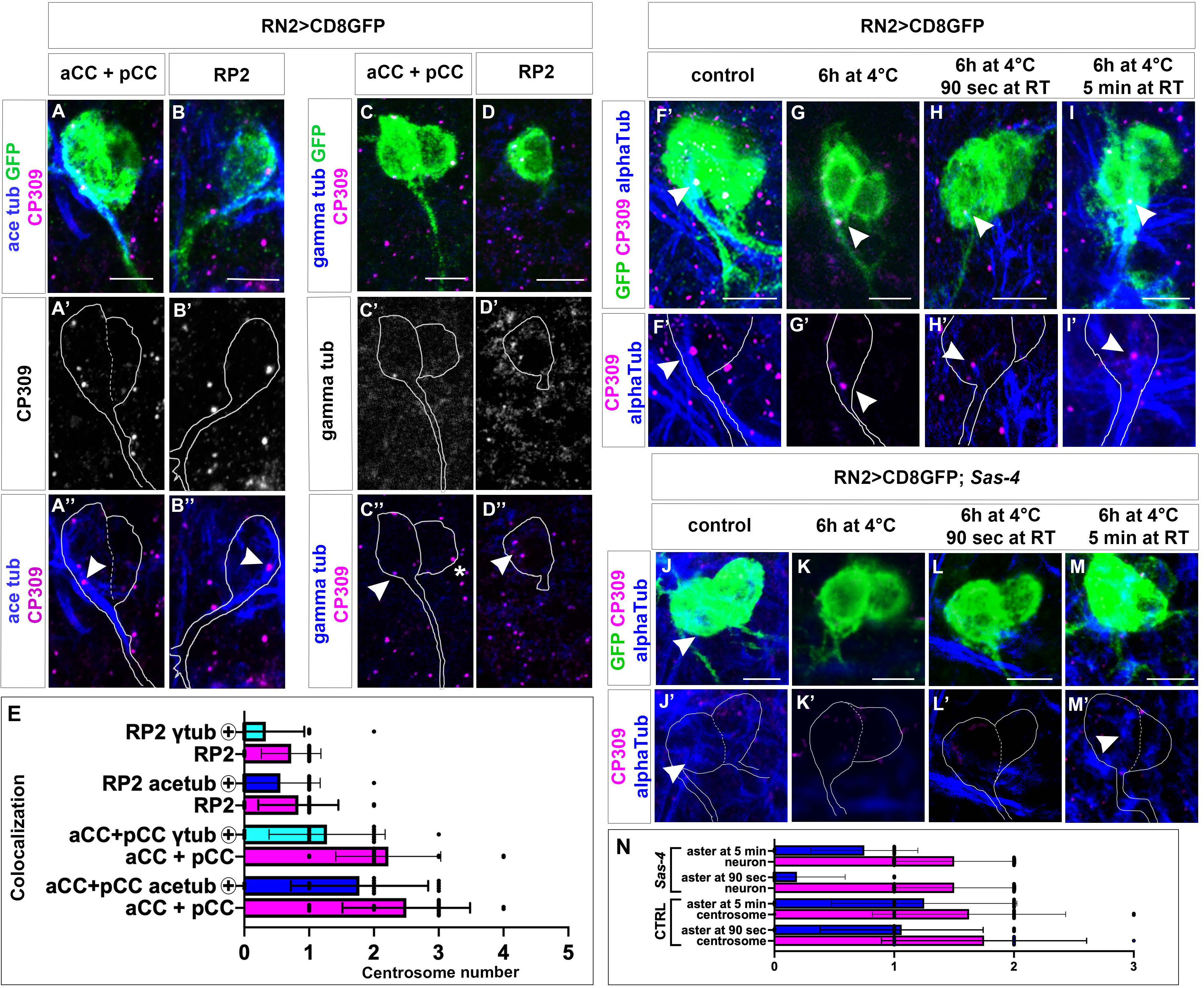
Pioneer motor neurons have active centrosomes. (A) RN2>CD8GFP with aCC+pCC and RP2 (B) marked in green, Plp (CP309) in magenta and ace-tub in blue; scale bar 5 µm. (A’, B’) Plp (CP309) marking all centrioles and the neuronal body drawn from the channel marking the neuron (A, B). (A’’, B’’) Plp (CP309) marking all centrioles and acetylated-tubulin antibody marking stable MTs with the neuronal body drawn from the channel marking the neuron (A, B). (C) aCC+pCC and RP2 (D) marked in green, Plp (CP309) in magenta and y-tub in blue. (C’, D’) Plp (CP309) marking all centrosomes and the neuronal body drawn from the channel marking the neuron (C, D). (C’’, D’’) Plp (CP309) marking all centrosomes and anti-acetylated-tubulin antibody marking stable MTs with the neuronal body drawn from the channel marking the neuron (C, D). (E) Quantification of the total number of centrosomes in aCC+pCC and RP2 neurons and quantification of the numbers of these which colocalize with y-tub (cyan column n=18 neurons) and acetylated tubulin (dark blue column, n=18 neurons). (F-I) *RN2>CD8GFP* with aCC+pCC marked in green, Plp (CP309) in magenta and alpha-tub in blue; scale bar 5 µm. (F’-I’) Plp (CP309) marking all centrosomes in magenta and anti-alpha-tubulin antibody marking all MTs in blue, with the neuronal body drawn from the channel marking the neuron (F-I). (F) Control embryos, 0h at 4°C; (G) embryos incubated 6h at 4°C; (H) embryos incubated 6h at 4°C and allowed to recover for 90 seconds at RT; (I) embryos incubated 6h at 4°C and allowed to recover for 5 min at RT. (J-M) *RN2>CD8GFP; Sas-4* mutant embryos with aCC+pCC marked in green, Plp (CP309) in magenta and alpha-tub in blue; scale bar 5 µm. (J’-M’) Plp (CP309) marking all centrosomes in magenta and anti-alpha-tubulin antibody marking all MTs in blue, with the neuronal body drawn from the channel marking the neuron (J-M). (J) Control embryos, 0h at 4°C; (K) embryos incubated 6h at 4°C; (L) embryos incubated 6h at 4°C and allowed to recover for 90 seconds at RT; (M) embryos incubated 6h at 4°C and allowed to recover for 5 min at RT. Scale bar 5 µm. (N) Quantification of the numbers of centrosomes in aCC+pCC and RP2 neurons and quantification of the MT asters at 90 secs and 5 min of reassembly time in control and *Sas-4* mutant neurons. In control neurons, MTs asters colocalize with the centrosome; n= 32 aCC and pCC and n=16 RP2.

To determine whether centrosomes function as active MTOCs in these neurons, we depolymerized microtubules and monitored their regrowth from the centrosome (Fig. 5F–I) (Brodu *et al*, 2010). This showed that microtubule regrowth initiates at the centrosome following depolymerization (Fig. 5 F–I). After 90 secs of recovery at RT we could observe an ⍺-tubulin aster emerging from a Plp positive centrosome in 60.7% of the cases and this increased to 76.9% after 5 min at RT (Fig. 5 N, n=32 neurons). Together, these results indicate that neuronal centrosomes are functionally active during the early stages of axon formation and serve as sites of microtubule organization in these cells.

### Sas-4 mutants without centrosomes reassemble MTs at a slower rate

Given that *Sas-4* mutant pioneer neurons lack centrosomes but are still able to extend axons, we asked whether the absence of centrosomes affects the efficiency or dynamics of MT reassembly. To address this question, we performed a MT disassembly and regrowth assay in *Sas-4* mutant embryos (Fig. 5 J-M and N). Following MT depolymerization, ⍺-tubulin-positive asters were observed emerging in aCC/pCC neurons, although with delayed kinetics compared with control neurons. After 90 sec of recovery at RT, ⍺-tubulin asters were detected in only 12.5% of neurons, increasing to 50% after 5 min of recovery (Fig. 5 L, M, N n=24 neurons). These findings are consistent with the notion that centrosomes are not essential for MT reassembly in pioneer neurons but may facilitate the efficiency and timing of MT nucleation during early axonal development.

### PNS neurons maintain active centrosomes

In *Drosophila* embryos, the dorsal cluster neurons project their axons in a highly organized manner to form the sensory axonal tracts (Ghysen *et al*, 1986). During development, the axons of these neurons extend dorsally and medially, connecting with their target regions and fasciculating with the intersegmental nerve (ISN) to form a bundled tract for efficient sensory signal routing (Hartenstein, 1988). Sensory neurons in *Drosophila* embryos are known to have centrioles. These centrioles are crucial for forming basal bodies in ciliated cells, including type-I sensory neurons, which are essential for the development of cilia (Jana *et al*, 2016). Since we detected axonal guidance phenotypes in the axons extending from the dorsal cluster, we tested for the presence of centrioles in dorsal cluster sensory neurons (Fig. 6A). We detected centrioles in the soma of most of these neurons and quantified the number of centriole-positive ddaC neurons (Fig. 6A’’, B, and C’, arrowhead), finding centrioles in 81.4% of the total (Fig. 6B, n=43). Importantly, we also detected centrioles in dorsal bipolar neuron (dbp, also known as dbd), the proposed pioneer neuron of the dorsal cluster (Fig. 6A’’ and C’, arrow) (Martin *et al*, 2008; Merritt & Whitington, 1995). In *Sas-4* embryos, the presence of centrioles in the dorsal cluster neurons was significantly lower (Fig. 6D). In many embryos, we could not detect any centrioles in the soma of the dorsal cluster neurons (Fig. 6E).

**Fig. 6.**
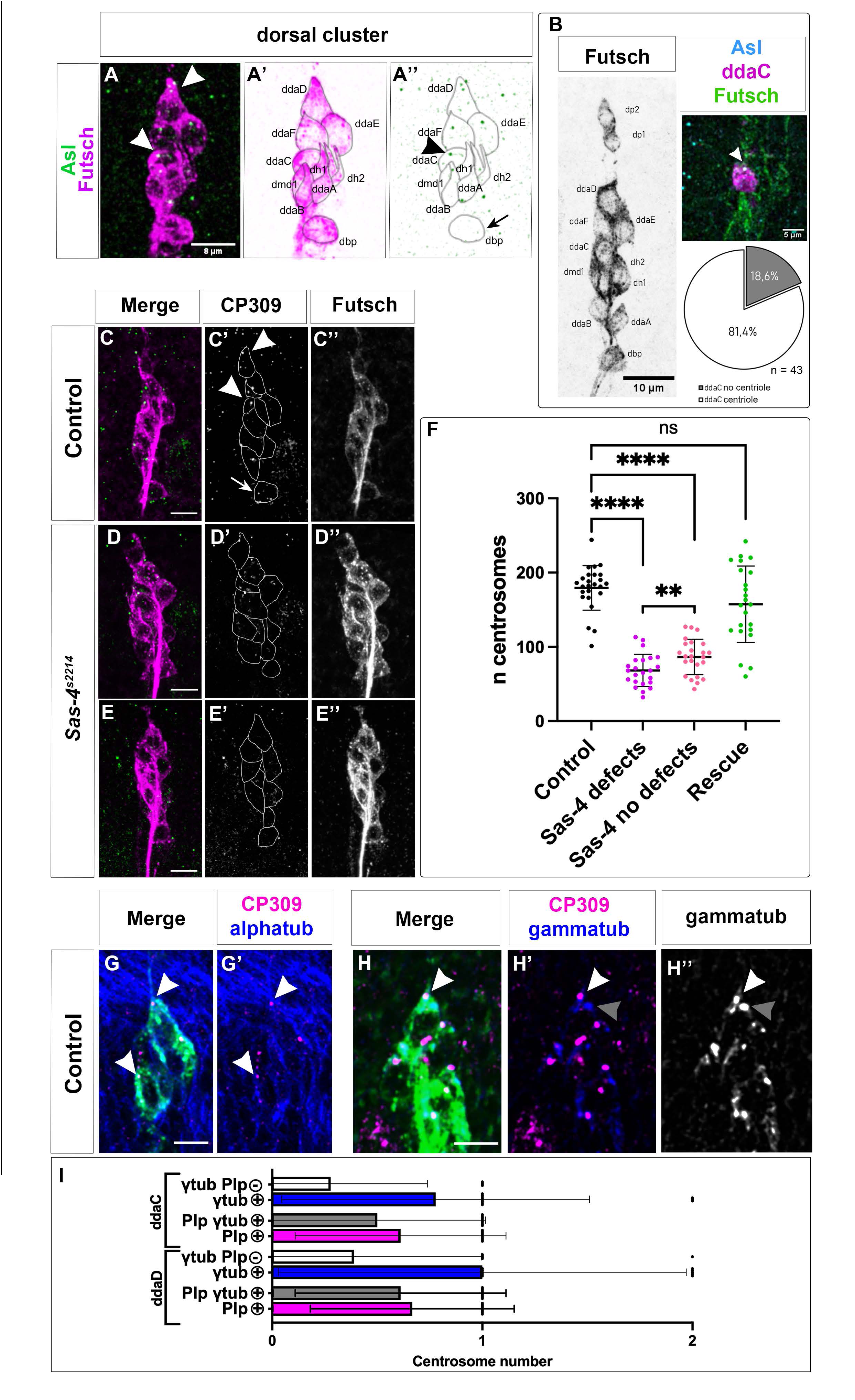
PNS neurons maintain active centrosomes. (A) Dorsal cluster neurons in a stage 15 embryo marked with anti-Futsch (22C10) in magenta to mark the sensory neurons and Asterless (Asl) in green to mark the centrosomes. (A’) Inverted channels with neurons identified and (A’’) cell bodies drawn to show the centrosomes within each neuron. Scale bar is 8 µm (B) Stage 16 dorsal cluster with the neurons identified and ddaC marked in green with ppkGAL4 driving UASCD8GFP marking class IV dendritic arborisation neurons, with centriole in cyan and Futsch in green. Scale bar is 5 µm. Quantification of the number of ddaC neurons with centrosomes (81.4% n=43). (C) Control and *Sas-4* (D,E) dorsal clusters marked with Futsch in magenta and Plp (CP309) in green. (C’, D’ and E’) Plp (CP309) marking all centrosomes and cell bodies drawn from the other channel (C’’, D’’ and E’’); scale bar Is 5 µm. (F) Quantification of the number of centrosomes per stage 16 embryonic segmental area in Control, *Sas-4/+* embryos, (mean 179.4 ± 30.1, n=25), *Sas-4* embryos displaying PNS mutant phenotypes (mean 68.2 ± 21.6, n=23), *Sas-4* embryos without detectable PNS mutant phenotypes (mean 86.3 ± 23.8, n=24) and *Sas-4* embryos rescued by ubiquitous expression of Sas-4 (mean 157.3 ± 51.45, n=23); unpaired t-test with Welch’s correction control vs *Sas-4* p<0.0001, control vs *Sas-4* no defects p<0.0001, control vs *Sas-4* rescue p=0.08 and *Sas-4* vs *Sas-4* no defects p=0.009. (G) Detail of the dorsal part of the dorsal cluster stained marked with Futsch (green), Plp (CP309) (magenta), and alpha-tubulin (blue); arrowheads point to the colocalization of centrosomes with MTs. (H) Detail of the dorsal part of the dorsal cluster stained marked with Futsch (green), Plp (CP309) (magenta), and γ-tubulin (blue); arrowheads point to the colocalization of centrosomes with γ-tubulin; scale bar is 2.5 µm. (I) Quantification of the number of centrosomes in ddaC and ddaD neurons which are positive for Plp (magenta column) and for Plp and γ-tubulin (grey column); and number of γ-tubulin foci (blue column) which are negative for Plp (white column); n=18 neurons.

Based on our findings in the CNS and PNS, we decided to quantify the total number of centrosomes in *Sas-4* embryos and compare them to controls. To do so, we selected equal segmental areas spanning the dorsal cluster and the lateral chordotonal organs (lch) and counted the total number of centrioles in stage 16 control embryos, *Sas-4* homozygous embryos displaying mutant PNS phenotypes, *Sas-4* homozygous embryos with no apparent mutant phenotype and *Sas-4* mutants rescued by ubiquitous Sas-4 expression (Fig. 6F). We observed that *Sas-4* mutants had approximately one-third of the centrioles found in the control group consistent with our previous findings (Fig. 1 and 2). Additionally, ubiquitous expression of Sas-4 in *Sas-4* mutant embryos restored centriole numbers to levels comparable to those of controls (Fig. 6F). These results are consistent with a progressive loss of centrioles during embryonic development in *Sas-4* mutant embryos. Notably, embryos lacking overt phenotypes retained, on average, slightly more centrosomes than embryos displaying mutant phenotypes, although centrosome numbers remained substantially lower than in wild-type embryos (Fig. 6F). Together, these observations suggest that centrosomes are not strictly required for neuronal development in this context but may contribute to its robustness or proper execution. To assess whether sensory neuron centrosomes are associated with MT nucleation machinery, we analysed their colocalization with α-tubulin and γ-tubulin. We found that sensory neuron centrosomes consistently colocalise with both markers (Fig. 6 G, H and I), indicating the presence of structural microtubules and nucleation components at the centrosome. We also analysed γ-tubulin foci that did not colocalize with detectable Plp (Fig. 6 H′, H″, grey arrows), as these are likely to correspond to other MTOCs present in these neurons. We could detect that about 40% of γ-tubulin foci in these PNS neurons co-localize with Plp (Fig. 6 I). Together, these observations suggest that sensory neuron centrosomes, similarly to those in motor neurons, remain active during the early stages of axon development, while additional γ-tubulin-positive, Plp-negative foci may reflect the presence of other MTOCs in these neurons.

### Centrosome loss induces motor axon fibre tortuosity

Axonal tortuosity (wiggliness) is a measure of the irregularities or abnormalities in the morphological structure of axons which can be associated with neurodegeneration (Bando *et al*, 2015). When axons are “wiggly,” it suggests that they are not following their usual straight or organized paths, possibly due to changes in axonal cytoskeletal structure or cell-adhesion, but not much is known about what triggers axonal wiggliness. Qualitative examination of confocal projections showed that aCC and RP2 axons are straighter in controls than in *Sas-4* mutants, revealing the tortuosity of these axons (Fig. 7 A-E, asterisks). Apart from the previously characterised phenotypes such as axonal crossings (Fig. 7 B, arrowhead) and misrouting, we could observe axonal wiggliness at the CNS exit points of *Sas-4* motor neurons (Fig. 7 B’, D and E, asterisks). Since we were interested in determining the proportion of wiggly axons in the *Sas-4* mutant and *Sas-4 RNAi* embryos, we quantified this for the aCC and RP2 neurons (Fig. 7 G). In *Sas-4* mutants we could detect an average of 23% aCC and 30% RP2 wiggly axons per total number of aCC and RP2 axons, which was significantly different from the proportion of wiggly axons in the controls (6% wiggly aCC and 8% wiggly RP2 axons). The same was observed for the Sas-4 downregulation in aCC, pCC and RP2 neurons (24% aCC and 25% RP2 wiggly axons per total number of aCC and RP2 axons). We also quantified axonal wiggliness via the ratio of Path Length in 2D (PL2D) normalized to Euclidean Distance in 2D (PL2D/ED2D) for each aCC and RP2 axonal path (Arshadi *et al*, 2021). We could detect significant differences between control and *Sas-4* mutant conditions (Fig. 7H). To correlate axonal wiggliness with the absence of centrosomes, we quantified the axonal morphology defects associated with centrosome loss. We scored aCC and RP2 axons as either tortuous (wiggly) or non-tortuous (Fig. 7 I-K). Neurons lacking centrosomes exhibited a substantially higher frequency of tortuous axons compared with neurons with detectable centrosomes (37/68 vs. 7/59, Fig. 7 L). Consistent with this observation, a chi-square test revealed a significant association between centrosome loss and axonal tortuosity (χ² = 23.41, df = 1, p < 0.0001).

**Fig. 7.**
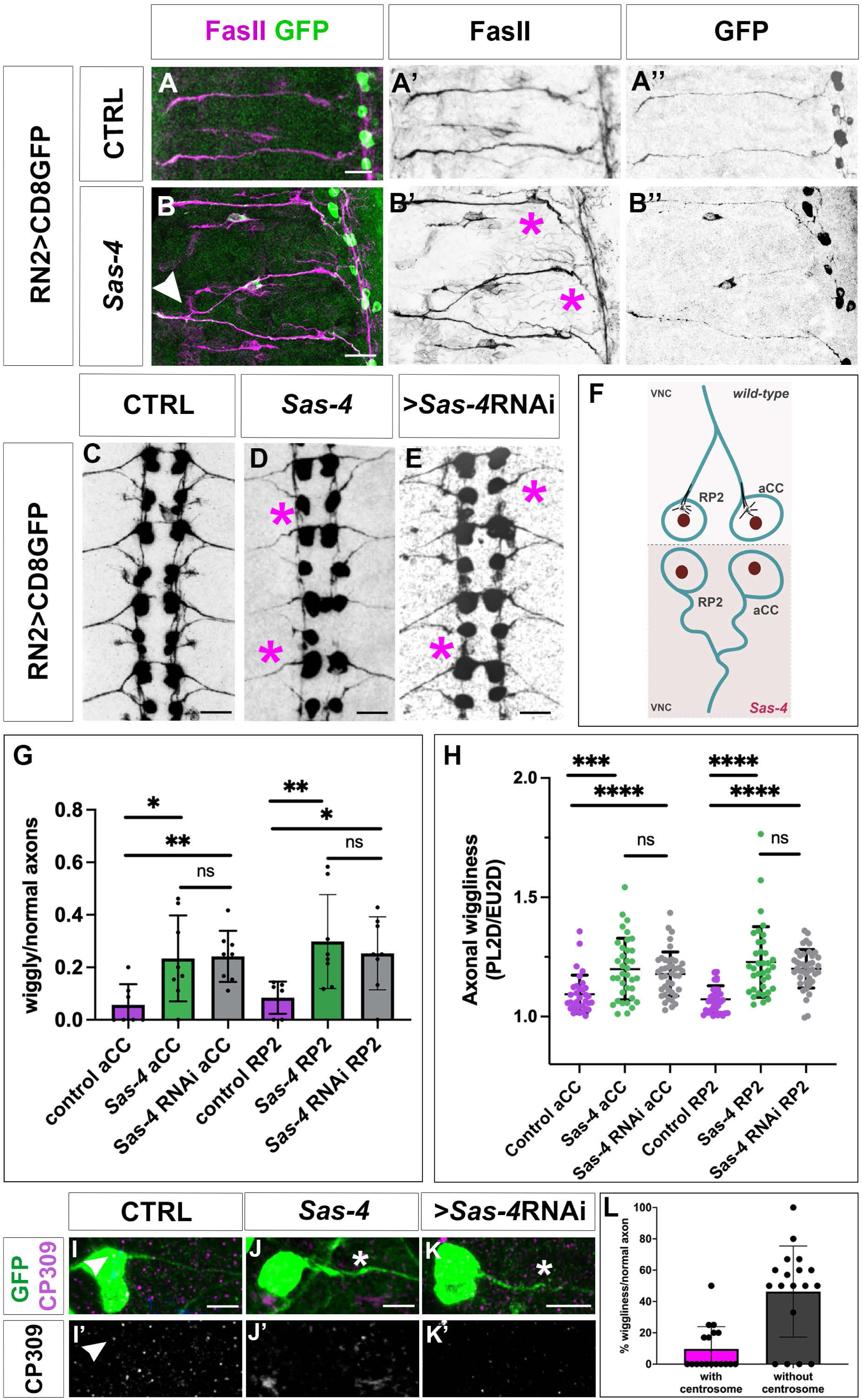
*Sas-4* loss of function induces wiggly axon phenotypes. (A-E) Confocal images of stage 16 RN2GAL4, UASCD8GFP embryonic lateral segments (A,B) or the ventral midline (CEF) in wt (A, C), *Sas-4* mutant background (B, D) or Sas-4 RNAi downregulation (E). (A’,B’) axons marked with anti-FasII and (A’’, B’’, C, D, E) axons marked with anti-GFP antibody, marking the axonal membranes. Scale bar is 10 µm in A-E. (F) Schematic representation of the wiggly axon phenotypes; in *Sas-4* aCC and RP2 neurons, the absence of active centrosomes gives rise to axonal tortuosity. (G) Quantification of the proportion (wiggly/normal axons) of wiggly axons in control, *Sas-4* mutant and Sas-4 RNAi downregulation for aCC and RP2 neurons; Mann-Whitney test p=0.025 in control vs *Sas-4* aCC (n=8 embryos); p= 0.0026 in control vs Sas-4 RNAi aCC (n=8 embryos) and p=0.0065 in control vs *Sas-4* RP2 (n=8 embryos); p=0.0149 in control vs Sas-4 RNAi RP2 (n=8 embryos). (H) Quantification of axonal wiggliness via the ratio of Path Length in 2D (PL2D) normalized to Euclidean Distance in 2D (PL2D/ED2D) for each aCC and RP2 axonal path. Mann-Whitney test p=0.0002 in control (n= 33 axons) vs *Sas-4* aCC (n=36 axons); p< 0.0001 in control vs Sas-4 RNAi aCC (n=41 axons) and p< 0.0001 in control (n=35 axons) vs *Sas-4* RP2 (n= 36 axons); p< 0.0001 in control vs Sas-4 RNAi RP2 (n= 41 axons). (I-K) Example of quantified neurons: confocal projections of stage 14 RN2GAL4, UASCD8GFP aCC neurons stained with GFP (green) and CP309 (magenta); (I) control with centrosome; (J) *Sas-4* mutant without centrosome; (K) Sas-4 downregulation without centrosome; scale bar 5µm. (L) Quantification of aCC and RP2 neurons with and without centrosomes and with or without tortuous axons.

It was previously shown increased JNK activity, can result in axonal wiggliness (Karkali *et al*, 2023). However, when we checked JNK signalling, we could not detect any increase in JNK activity in *Sas-4* neurons (Supl. Fig. 4). In conclusion, although CL does not completely prevent axonal extension, it is associated with morphological abnormalities in axonal architecture (Fig. 7F), consistent with broader alterations in cytoskeletal organization and suggesting a contribution of centrosomes to normal axonal growth.

## Discussion

With this work, we provide evidence supporting a role for centrosomes in axonal extension, axon guidance, and muscle development during embryogenesis in vivo. Notably, centrosome loss does not abolish axonal extension but is associated with defects in axonal architecture (Fig. 7F), consistent with broader perturbations of cytoskeletal organization and suggesting that centrosomes contribute to normal axonal growth. Previous studies have shown that *Sas-4* and *Sas-6* mutant flies can reach adulthood (Basto *et al*., 2006; Fatalska *et al*., 2021; Peel *et al*., 2007), which initially seems incompatible with an important role for the centrosome during embryonic development. However, we found that *Sas-4* and *Sas-6* homozygous null conditions result in 50% of embryonic lethality.

*Sas-4* and *Sas-6* LOF embryos display variable motor axon phenotypes (Figs. 1 and 3), consistent with a role for centrosomes in axonal extension. Additionally, Sas-4 and Sas-6 LOF embryos exhibit muscle phenotypes (Figs. 2 and 3), consistent with a contribution of centrosomes to muscle development. Hence, the embryonic axon guidance phenotypes have a dual contribution: non-autonomous one from muscle CL (Fig. 8A) and an autonomous one from axonal CL (Fig. 8B). Since innervation does not influence the patterning, morphogenesis, maintenance, or physiological development of somatic muscles in *Drosophila* embryos, the muscle defects observed are most likely due to CL in muscle cells (Broadie & Bate, 1993). CL in muscle cells may impair cilia formation and nuclear positioning, two processes important for muscle fibre development (Loh *et al*, 2023; Metzger *et al*, 2012; Tillery *et al*, 2018). Expressing *Sas-4 RNAi* under the control of the muscle-specific *Mef2-GAL4* driver induced penetrant muscle phenotypes, consistent with previous results reporting the lethality of these embryos (Schnorrer *et al*, 2010). *Sas-6* embryos exhibited milder muscle phenotypes than *Sas-4*, aligning with previous findings that driving *Sas-6 RNAi* with a Mef2-GAL4 driver induced adult pharate lethality (Schnorrer *et al*., 2010).

**Fig. 8.**
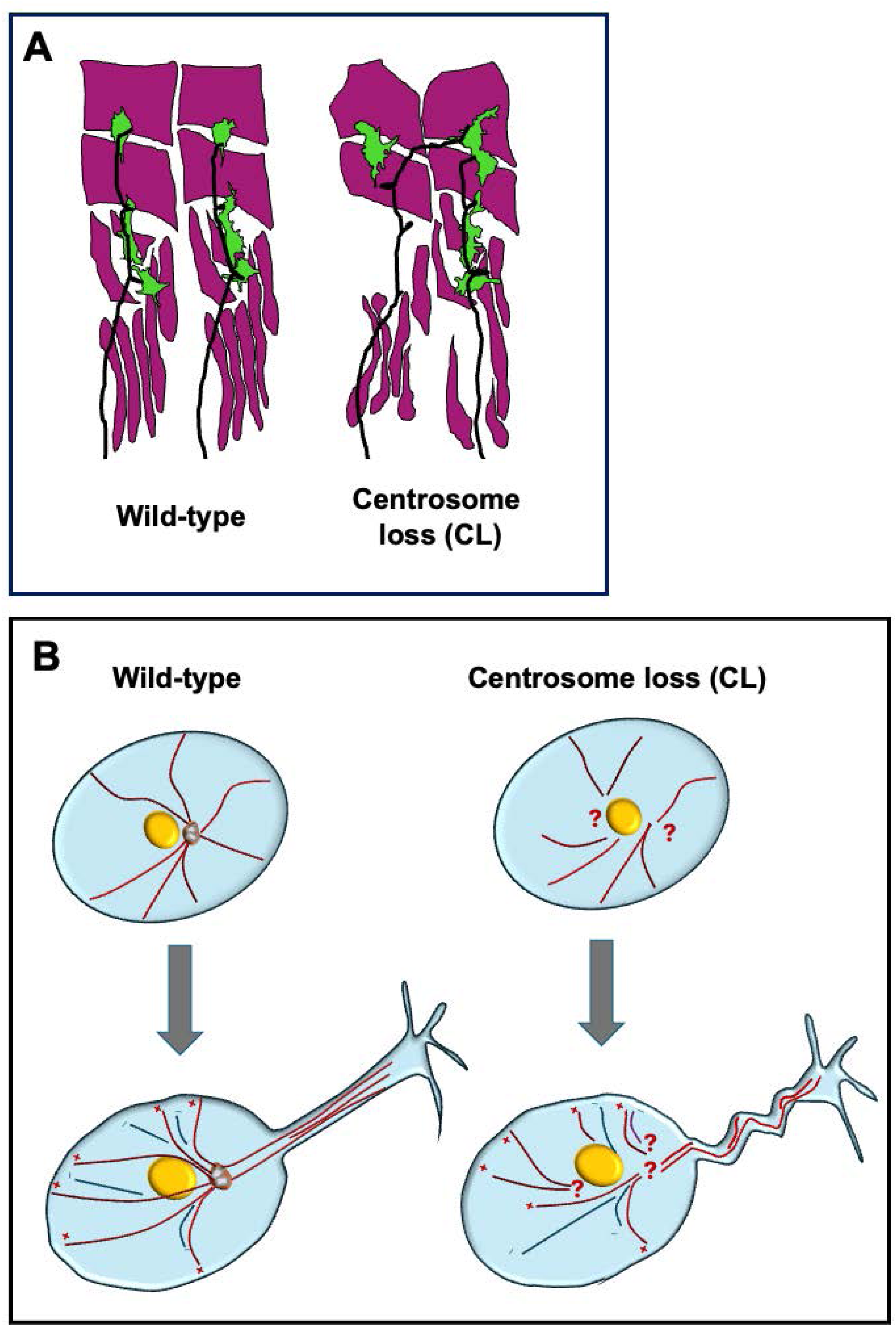
Centrosome loss affects axonal morphogenesis and muscle development. Schematic representations of CL induced axonal phenotypes. CL disrupts axonal morphogenesis and pathfinding through two distinct mechanisms. (A) First, in a neuron non-autonomous manner, CL impairs muscle development, which in turn leads to axon guidance defects. (B) Second, in a neuron-autonomous manner, CL compromises the integrity of the MT cytoskeleton, resulting in defective axon pathfinding and increased axonal tortuosity. Taken together, these observations suggest that the axonal phenotypes associated with centrosome loss reflect the combined impact of altered muscle development and changes in axonal architecture, consistent with a contribution of centrosomes to multiple processes involved in axon development.

What could explain the variability in the consequences of CL during embryonic development? Many disease-causing mutations exhibit mild or no effects in some individuals, a phenomenon known as incomplete phenotype penetrance. This phenomenon is still poorly understood, but recent studies in *Drosophila* suggest it is stochastic, akin to flipping a coin (Marmion *et al*, 2023). Although the reported study focuses on fluctuations in signalling, a similar scenario could be imagined for centrosomes. In cases of centrosome duplication impairment which induce CL, distribution would vary among dividing cells as development proceeds. Early embryonic mitotic cycles can partly explain the differences in phenotype expressivity. The first 13 mitotic cycles of *Drosophila* embryos are rapid, synchronous, and occur without cell division. Subsequently, three rounds of patterned mitosis occur (mitoses 14, 15, and 16). Following mitosis 16, many cells enter their first G1 phase (Edgar & O’Farrell, 1990). However, the nervous system lineage does not enter a terminal interphase after mitosis 16 and continues to divide according to an independent program (Bodmer & Jan, 1987; Hartenstein *et al*, 1987). Similarly, muscle cells, continue dividing until embryonic stage 10 (Nabel-Rosen *et al*, 2005). These additional cell divisions may account, at least in part, for the appearance and variable severity of the observed phenotypes, as the consequences of centrosome loss are likely to become more pronounced with successive rounds of cell division owing to progressive depletion of the maternal contribution. In agreement with this, when we quantified the number of centrioles present in *Sas-4* embryos displaying axonal phenotypes, we could detect a decrease of about 1/3 (Fig. 6), with consistent results when we quantified the numbers of centrioles in *Sas-4* aCC, pCC and RP2 neurons (Fig. 4)

### Centrosome localization in neurons and implications for axonal morphology

Here we show that centrosomes are present in motor and sensory neurons and that, in motor neurons, these centrosomes colocalize with acetylated tubulin and γ-tubulin and participate in MT polymerization, consistent with their functioning as active MTOCs in these cells (Figs. 4, 5 and 6). A previous study failed to detect colocalization of centrosomes with γ -tubulin in neurons (Nguyen *et al*., 2011). However, this study used Fizzy-related-GFP (Fzr-GFP) to visualise centrioles and, while it is clear Fzr is localized to centrioles during the cell cycle (Meghini *et al*, 2016), it is not clear this localization is maintained in all fully differentiated cells (Djabrayan *et al*, 2014; Huynh *et al*, 2009). This, or differences in microscopy approaches, may account for the discrepancy between our findings and previous reports. Furthermore, centrosome localization and MT-organizing activity may be particularly important during the initial stages of axon outgrowth (embryonic stages 12-13), rather than after axonal projections have already formed (embryonic stages 15-16), when many of these earlier observations were made.

The persistence of centrosome-associated phenotypes in sensory neurons is particularly interesting given that we detect additional non-centrosomal MTOCs. The sensory neurons analysed here belong to the dendritic arborization (da) neuron class and subsequently develop extensive and highly branched dendritic arbours. The presence of additional acentrosomal MTOCs may therefore be related to their more complex morphology and the need to organise microtubules throughout elaborate dendritic compartments. Indeed, non-centrosomal sites of MT nucleation have been described in developing sensory neurons and are thought to contribute to dendritic microtubule organisation (Ori-Mckenney *et al*, 2012). Nevertheless, CL remained associated with neuronal phenotypes in these cells, suggesting that alternative MTOCs may not fully compensate for centrosome function during early stages of neuronal development. In contrast, motor neurons, which develop a comparatively simpler dendritic architecture, may depend more strongly on centrosome-based MT organisation during early axon outgrowth.

Regarding centrosome localization and activity in neurons, we show that axonal morphology in embryos with fewer centrosomes is altered. To our knowledge, axonal tortuosity represents the first morphological axon phenotype reported in association with centrosome loss in *Drosophila*. Under conditions of CL, we observe tortuous (wiggly) axons, suggesting that while CL does not prevent neurons from extending axons, it does affect axonal outgrowth. CL could induce this phenotype by disrupting the neuronal cytoskeleton and preventing MTs to form, be severed and transported into the axons. It is known that alterations in the neuronal cytoskeleton, caused by signalling deregulation such as increased JNK activity, can result in axonal wiggliness (Karkali *et al*., 2023). Also, alterations in MT dynamics in mammalian neurons have been shown to decrease axonal elongation, but not its formation, giving rise to wiggly axons (Rochlin *et al*, 1996). In addition, axonal tortuosity is one of the hallmarks of neurodegeneration (Salvadores *et al*, 2020), which makes this phenotype a good entrance point for the analysis of the progression of various neurodegenerative diseases.

The loss of centrosome function has been documented to disrupt various developmental stages in vertebrates. In humans, CL is associated with hereditary spastic paraplegia, Bardet-Biedl syndrome, the formation of cystic kidneys, disrupted left-right asymmetry, and microcephaly (Badano *et al*, 2005; Jaiswal & Singh, 2021). Demonstrating that centrosome loss in *Drosophila* affects the development of the nervous and muscular systems highlights the potential of this model organism for investigating the mechanisms underlying related human diseases.

## Acknowledgments

We are grateful to M. Llimargas and J. Casanova for generously providing access to their laboratory for specific aspects of this work and for their continuous support throughout the study. We thank B. Estrada, C. González, K. Karkali, C. Klämbt, E. Martin-Blanco, J. Raff, S. Rumpf, Y. Zheng, the Bloomington Drosophila Stock Center (BDSC) and the Vienna Drosophila Stock Center (VDRC) for fly stocks and reagents. We thank K. Karkali for valuable assistance with specific quantifications and technical procedures. We extend our gratitude to Z. Novak and J. Raff for sharing two unpublished CRISPR-Cas9 made *Sas-6* fly stocks with us. Thanks also go to R. Mendez for technical assistance in all the experiments and M. Bosch for assistance and advice with confocal microscopy and software. B.G. is the recipient of a Beatriu de Pinós post-doctoral fellowship from the AGAUR (2022BP00125). This work was supported by a grant from the Spanish *Ministerio de Ciencia y Innovación* (PID2021-125860NB-I00), the Universitat de Barcelona, and a grant from the Generalitat de Catalunya (2021 SGR 1455).

## Author contributions

Conceptualization: S.J.A. Methodology and Experiments: B.G., J.P., J.S.-A., J.C.-R. and S.J.A. Analysis: B.G., J.P., J.S.-A. and S.J.A. Writing: S.J.A. Reviewing and editing: S.J.A. and B.G. Supervision and Funding: S.J.A.

---

**Sup. Fig 1.**
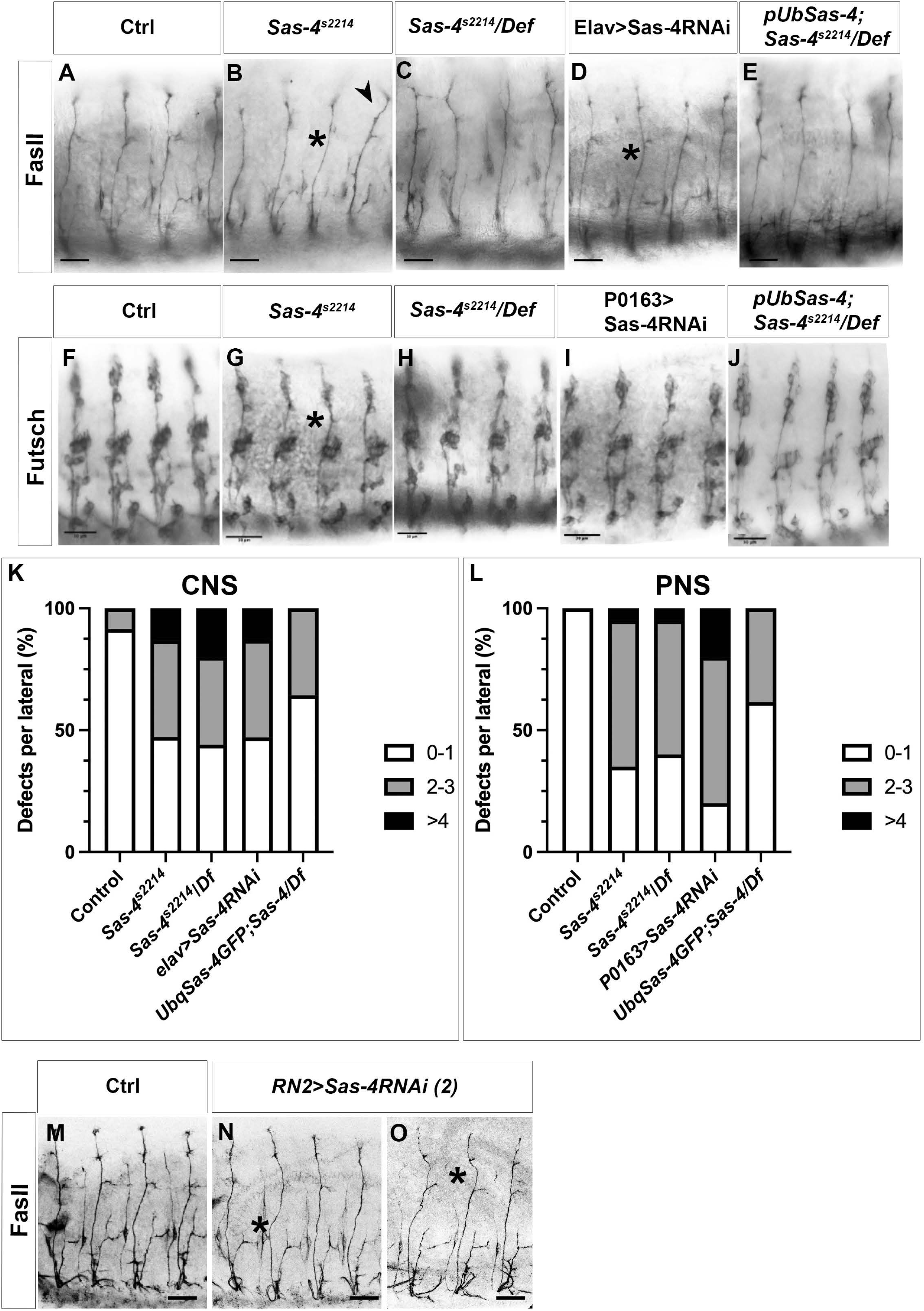

**Sup. Fig 2.**
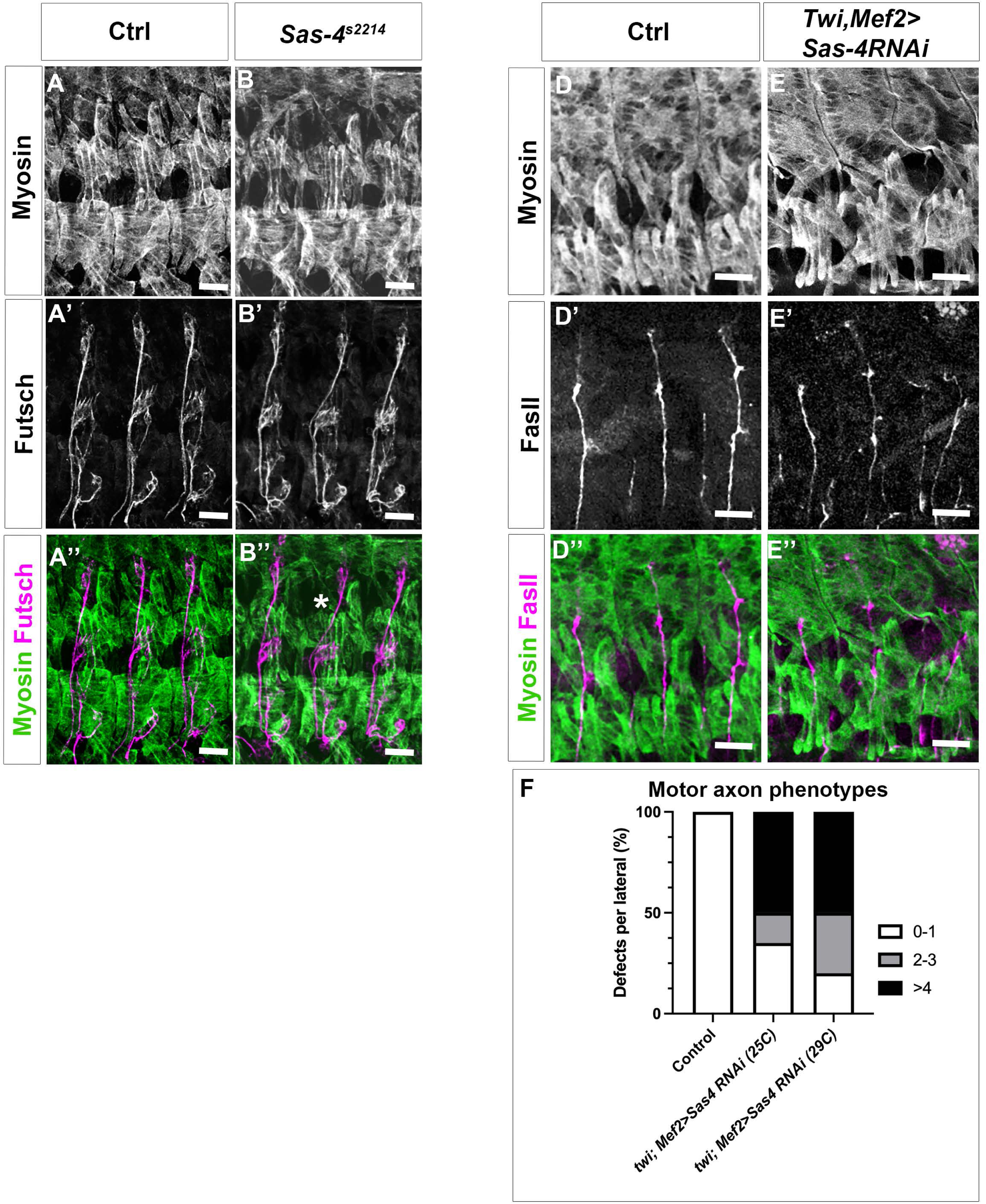

**Sup. Fig 3.**
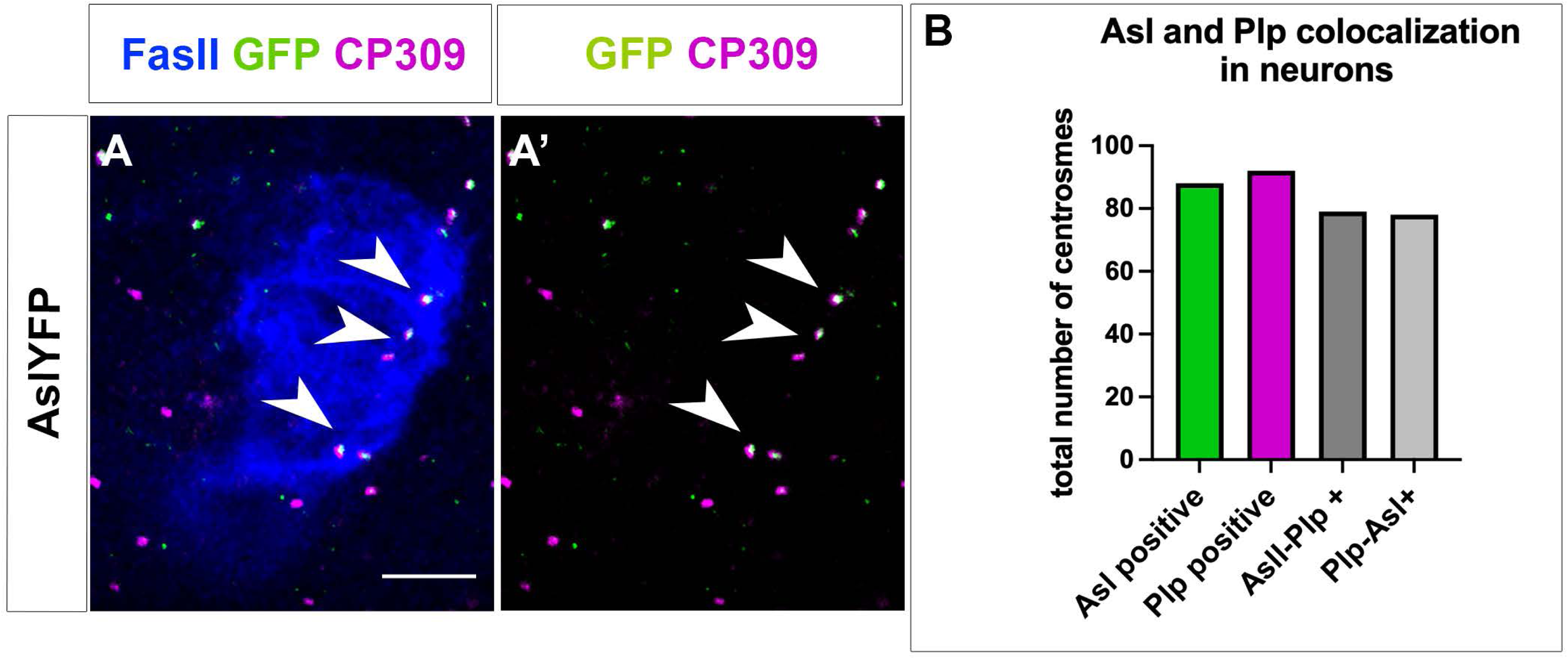

**Sup. Fig 4.**
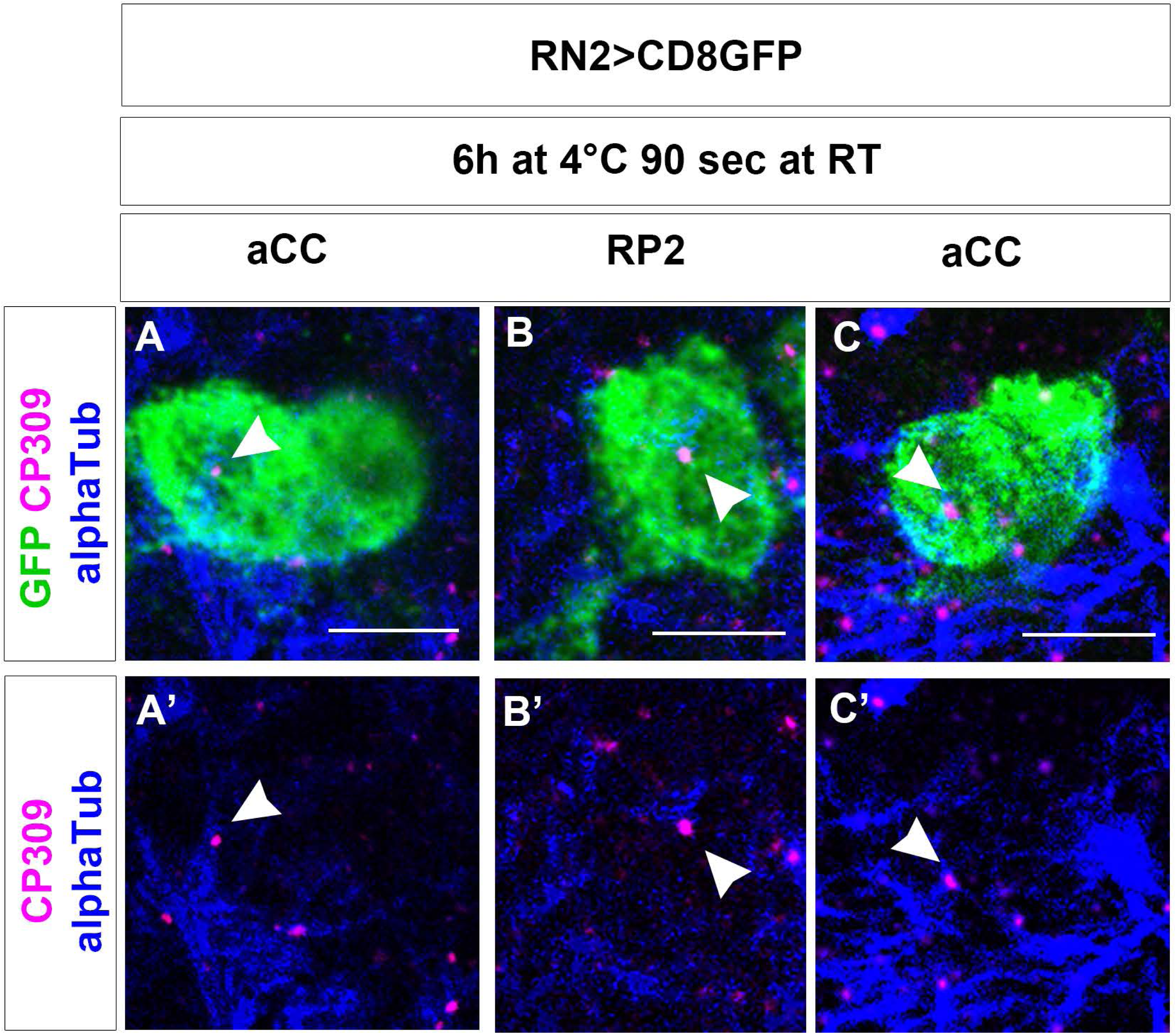

**Sup. Fig 5.**
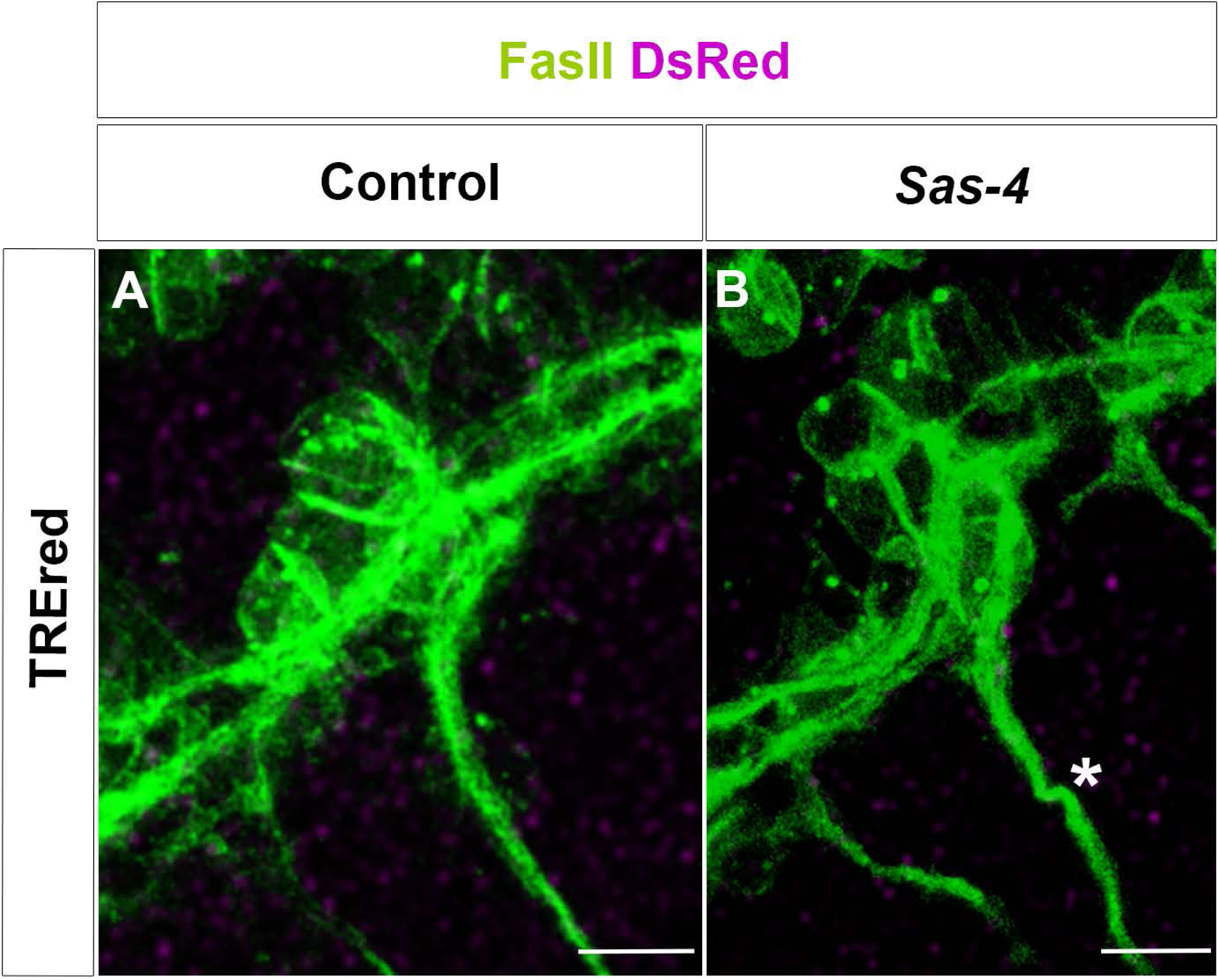

## Notes

### Competing Interest Statement

The authors have declared no competing interest.

### Summary of Updates

We have updated the text and some of the figures.

## References

Ahmad FJ, Baas PW (1995) Microtubules released from the neuronal centrosome are transported into the axon. Journal of Cell Science 108 ( Pt 8): 2761–2769

Anda FCd, Meletis K, Ge X, Rei D, Tsai L-H (2010) Centrosome Motility Is Essential for Initial Axon Formation in the Neocortex. The Journal of Neuroscience 30: 10391–10406

Anda FCd, Pollarolo G, Silva JSD, Camoletto PG, Feiguin F, Dotti CG (2005) Centrosome localization determines neuronal polarity. Nature Cell Biology 436: 704–708

Araújo SJ (2019) The Golgi Apparatus and Centriole, Functions, Interactions and Role in Disease - Centrosomes in Branching Morphogenesis. Results Problems Cell Differ 67: 323–336

Arshadi C, Günther U, Eddison M, Harrington KIS, Ferreira TA (2021) SNT: a unifying toolbox for quantification of neuronal anatomy. Nature Methods 18: 374–377

Badano JL, Teslovich TM, Katsanis N (2005) The centrosome in human genetic disease. Nature Reviews Genetics 6: 194–205

Bando Y, Nomura T, Bochimoto H, Murakami K, Tanaka T, Watanabe T, Yoshida S (2015) Abnormal morphology of myelin and axon pathology in murine models of multiple sclerosis. Neurochemistry International 81: 16–27

Basto R, Lau J, Vinogradova T, Gardiol A, Woods CG, Khodjakov A, Raff JW (2006) Flies without centrioles. Cell 125: 1375–1386

Bazzi H, Anderson KV (2014) Acentriolar mitosis activates a p53-dependent apoptosis pathway in the mouse embryo. Proc Natl Acad Sci U S A 111: E1491–1500

Bettencourt-Dias M, Glover DM (2007) Centrosome biogenesis and function: centrosomics brings new understanding. Nat Rev Mol Cell Biol 8: 451–463

Bodmer R, Jan YN (1987) Morphological differentiation of the embryonic peripheral neurons in Drosophila. Roux’s Arch Dev Biol: 69–77

Broadie K, Bate M (1993) Muscle development is independent of innervation during Drosophila embryogenesis. Development 119: 533–543

Brodu V, Baffet AD, Droguen P-ML, Casanova J, Guichet A (2010) A developmentally regulated two-step process generates a noncentrosomal microtubule network in Drosophila tracheal cells. Developmental Cell 18: 790–801

Castellanos E, Dominguez P, Gonzalez C (2008) Centrosome Dysfunction in Drosophila Neural Stem Cells Causes Tumors that Are Not Due to Genome Instability. Current Biology 18: 1209–1214

Chang W, Antoku S, Östlund C, Worman HJ, Gundersen GG (2015) Linker of nucleoskeleton and cytoskeleton (LINC) complex-mediated actin-dependent nuclear positioning orients centrosomes in migrating myoblasts. Nucleus 6: 77–88

Cheng T, Agwu C, Shim K, Wang B, Jain S, Mahjoub MR (2023) Aberrant centrosome biogenesis disrupts nephron and collecting duct progenitor growth and fate resulting in fibrocystic kidney disease. Development 150

Conduit PT, Wainman A, Raff JW (2015) Centrosome function and assembly in animal cells. Nature Reviews Molecular Cell Biology 16: 611–624

del Castillo U, Lu W, Winding M, Lakonishok M, Gelfand VI (2015) Pavarotti/MKLP1 regulates microtubule sliding and neurite outgrowth in Drosophila neurons. Curr Biol 25: 200–205

Djabrayan Nareg JV, Cruz J, de Miguel C, Franch-Marro X, Casanova J (2014) Specification of Differentiated Adult Progenitors via Inhibition of Endocycle Entry in the *Drosophila Trachea*. Cell Reports 9: 859-865

Edgar BA, O’Farrell PH (1990) The three postblastoderm cell cycles of Drosophila embryogenesis are regulated in G2 by string. Cell 62: 469–480

Fatalska A, Stepinac E, Richter M, Kovacs L, Pietras Z, Puchinger M, Dong G, Dadlez M, Glover DM (2021) The dimeric Golgi protein Gorab binds to Sas6 as a monomer to mediate centriole duplication. Elife 10

Fujioka M, Lear BC, Landgraf M, Yusibova GL, Zhou J, Riley KM, Patel NH, Jaynes JB (2003) Even-skipped, acting as a repressor, regulates axonal projections in Drosophila. Development 130: 5385–5400

Gartenmann L, Vicente CC, Wainman A, Novak ZA, Sieber B, Richens JH, Raff JW (2020) Drosophila Sas-6, Ana2 and Sas-4 self-organise into macromolecular structures that can be used to probe centriole and centrosome assembly. Journal of Cell Science 133: jcs244574

Ghysen A, Dambly-Chaudière C, Aceves E, Jan L, Jan Y (1986) Sensory neurons and peripheral pathways in Drosophila embryos. Rouxs Arch Dev Biol 195: 281–289

Gogendeau D, Basto R (2010) Centrioles in flies: The exception to the rule? Semin Cell Dev Biol 21: 163–173

Gomes ER, Jani S, Gundersen GG (2005) Nuclear Movement Regulated by Cdc42, MRCK, Myosin, and Actin Flow Establishes MTOC Polarization in Migrating Cells. Cell 121: 451–463

Goundiam O, Basto R (2020) Centrosomes in disease: how the same music can sound so different? Current opinion in structural biology 66: 74–82

Grzonka M, Bazzi H (2024) Mouse SAS-6 is required for centriole formation in embryos and integrity in embryonic stem cells. eLife 13: e94694

Harrison RG (1910) The outgrowth of the nerve fiber as a mode of protoplasmic movement. Journal of Experimental Zoology 9: 787–846

Hartenstein V (1988) Development of Drosophila larval sensory organs: spatiotemporal pattern of sensory neurones, peripheral axonal pathways and sensilla differentiation. Development 102: 869–886

Hartenstein V, Rudloff E, Campos-Ortega JA (1987) The pattern of proliferation of the neuroblasts in the wild-type embryo of Drosophila melanogaster. Rouxs Arch Dev Biol 196: 473–485

Hummel T, Krukkert K, Roos J, Davis G, Klämbt C (2000) Drosophila Futsch/22C10 Is a MAP1B-like Protein Required for Dendritic and Axonal Development. Neuron 26: 357–370

Huynh MA, Stegmüller J, Litterman N, Bonni A (2009) Regulation of Cdh1-APC function in axon growth by Cdh1 phosphorylation. J Neurosci 29: 4322-4327

Jacobs J, Goodman C (1989) Embryonic development of axon pathways in the Drosophila CNS. II. Behavior of pioneer growth cones. The Journal of Neuroscience 9: 2412–2422

Jaiswal S, Singh P (2021) Centrosome dysfunction in human diseases. Semin Cell Dev Biol 110: 113-122

Jana SC, Bettencourt-Dias M, Durand B, Megraw TL (2016) Drosophila melanogaster as a model for basal body research. Cilia: 1–7

Jeong S (2021) Molecular Mechanisms Underlying Motor Axon Guidance in Drosophila. Molecules and Cells 44: 549–556

Karkali K, Vernon SW, Baines RA, Panayotou G, Martín-Blanco E (2023) Puckered and JNK signaling in pioneer neurons coordinates the motor activity of the Drosophila embryo. Nature Communications 14: 8186

Kopeć S (1928) On the Influence of Intermittent Starvation on the Longevity of the Imaginal Stage of Drosophila Melanogaster. Journal of Experimental Biology 5: 204–211

Kushner EJ, Ferro LS, Liu J-Y, Durrant JR, Rogers SL, Dudley AC, Bautch VL (2014) Excess centrosomes disrupt endothelial cell migration via centrosome scattering. J Cell Biol 206: 257–272

Leask A, Obrietan K, Stearns T (1997) Synaptically coupled central nervous system neurons lack centrosomal γ-tubulin. Neuroscience Letters 229: 17–20

Leidel S, Gönczy P (2003) SAS-4 is essential for centrosome duplication in C elegans and is recruited to daughter centrioles once per cell cycle. Dev Cell 4: 431–439

Lin T-c, Neuner A, Schiebel E (2015) Targeting of γ-tubulin complexes to microtubule organizing centers: conservation and divergence. Trends in Cell Biology 25: 296–307

Lindhout FW, Portegies S, Kooistra R, Herstel LJ, Stucchi R, Hummel JJA, Scheefhals N, Katrukha EA, Altelaar M, MacGillavry HD et al (2021) Centrosome-mediated microtubule remodeling during axon formation in human iPSC-derived neurons. The EMBO Journal 40: e106798

Loh M, Bernard F, Guichet A (2023) Kinesin-1 promotes centrosome clustering and nuclear migration in the Drosophila oocyte. Development 150

Marmion RA, Simpkins AG, Barrett LA, Denberg DW, Zusman S, Schottenfeld-Roames J, Schüpbach T, Shvartsman SY (2023) Stochastic phenotypes in RAS-dependent developmental diseases. Current Biology 33: 807–816.e804

Marthiens V, Rujano MA, Pennetier C, Tessier S, Paul-Gilloteaux P, Basto R (2013) Centrosome amplification causes microcephaly. Nat Cell Biol 15: 731–740

Martin V, Mrkusich E, Steinel MC, Rice J, Merritt DJ, Whitington PM (2008) The L1-type cell adhesion molecule Neuroglian is necessary for maintenance of sensory axon advance in the Drosophila embryo. Neural Dev 3: 10

Meghini F, Martins T, Tait X, Fujimitsu K, Yamano H, Glover DM, Kimata Y (2016) Targeting of Fzr/Cdh1 for timely activation of the APC/C at the centrosome during mitotic exit. Nat Commun 7: 12607

Meka DP, Kobler O, Hong S, Friedrich CM, Wuesthoff S, Henis M, Schwanke B, Krisp C, Schmuelling N, Rueter R et al (2022) Centrosome-dependent microtubule modifications set the conditions for axon formation. Cell Reports 39: 110686

Merritt DJ, Whitington PM (1995) Central projections of sensory neurons in the Drosophila embryo correlate with sensory modality, soma position, and proneural gene function. J Neurosci 15: 1755–1767

Metzger T, Gache V, Xu M, Cadot B, Folker ES, Richardson BE, Gomes ER, Baylies MK (2012) MAP and kinesin-dependent nuclear positioning is required for skeletal muscle function. Nature 484: 120–124

Meyer-Gerards C, Bazzi H (2025) Developmental and tissue-specific roles of mammalian centrosomes. FEBS J 292: 709–726

Nabel-Rosen H, Toledano-Katchalski H, Volohonsky G, Volk T (2005) Cell Divisions in the Drosophila Embryonic Mesoderm Are Repressed via Posttranscriptional Regulation of string/cdc25 by HOW. Current Biology 15: 295–302

Nguyen MM, Stone MC, Rolls MM (2011) Microtubules are organized independently of the centrosome in Drosophila neurons. Neural Development 6: 38

Ori-Mckenney Kassandra M, Jan Lily Y, Jan Y-N (2012) Golgi Outposts Shape Dendrite Morphology by Functioning as Sites of Acentrosomal Microtubule Nucleation in Neurons. Neuron 76: 921–930

Pearl R, Parker SL (1921) Experimental Studies on the Duration of Life. I. Introductory Discussion of the Duration of Life in Drosophila. Am Nat 55: 481–509

Peel N, Stevens NR, Basto R, Raff JW (2007) Overexpressing Centriole-Replication Proteins In Vivo Induces Centriole Overduplication and De Novo Formation. Current Biology 17: 834–843

Poulton John S, Cuningham John C, Peifer M (2014) Acentrosomal Drosophila Epithelial Cells Exhibit Abnormal Cell Division, Leading to Cell Death and Compensatory Proliferation. Developmental Cell 30: 731–745

Poulton JS, Cuningham JC, Peifer M (2017) Centrosome and spindle assembly checkpoint loss leads to neural apoptosis and reduced brain size. Journal of Cell Biology 216: 1255–1265

Ricolo D, Castro-Ribera J, Araújo SJ (2021) Cytoskeletal players in single-cell branching morphogenesis. Developmental Biology 477: 22–34

Ricolo D, Deligiannaki M, Casanova J, Araújo SJ (2016) Centrosome Amplification Increases Single-Cell Branching in Post-mitotic Cells. Curr Biol 26: 2805–2813

Rochlin MW, Wickline KM, Bridgman PC (1996) Microtubule Stability Decreases Axon Elongation but Not Axoplasm Production. The Journal of Neuroscience 16: 3236–3246

Salvadores N, Gerónimo-Olvera C, Court FA (2020) Axonal Degeneration in AD: The Contribution of Aβ and Tau. Front Aging Neurosci 12: 581767

Sanchez-Huertas C, Freixo F, Viais R, Lacasa C, Soriano E, Luders J (2016) Non-centrosomal nucleation mediated by augmin organizes microtubules in post-mitotic neurons and controls axonal microtubule polarity. Nat Commun 7: 12187

Sánchez-Soriano N, Prokop A (2005) The influence of pioneer neurons on a growing motor nerve in Drosophila requires the neural cell adhesion molecule homolog FasciclinII. J Neurosci 25: 78–87

Schnorrer F, Schönbauer C, Langer CCH, Dietzl G, Novatchkova M, Schernhuber K, Fellner M, Azaryan A, Radolf M, Stark A et al (2010) Systematic genetic analysis of muscle morphogenesis and function in Drosophila. Nature 464: 287–291

Stevens NR, Raposo AASF, Basto R, Johnston DS, Raff JW (2007) From Stem Cell to Embryo without Centrioles. 17: 1498–1503

Stiess M, Maghelli N, Kapitein LC, Gomis-Rüth S, Wilsch-Bräuninger M, Hoogenraad CC, Tolić-Nørrelykke IM, Bradke F (2010) Axon extension occurs independently of centrosomal microtubule nucleation. Science 327: 704–707

Tillery MML, Blake-Hedges C, Zheng Y, Buchwalter RA, Megraw TL (2018) Centrosomal and Non-Centrosomal Microtubule-Organizing Centers (MTOCs) in Drosophila melanogaster. Cells 7

Varmark H, Llamazares S, Rebollo E, Lange B, Reina J, Schwarz H, Gonzalez C (2007) Asterless is a centriolar protein required for centrosome function and embryo development in Drosophila. Curr Biol 17: 1735–1745

Vinopal S, Dupraz S, Alfadil E, Pietralla T, Bendre S, Stiess M, Falk S, Ortega GC, Maghelli N, Tolić IM et al (2023) Centrosomal microtubule nucleation regulates radial migration of projection neurons independently of polarization in the developing brain. Neuron 111: 1241–1263.e1216

Younossi-Hartenstein A, Hartenstein V (1993) The role of the tracheae and musculature during pathfinding of Drosophila embryonic sensory axons. Dev Biol 158: 430–447

Yu W, Centonze VE, Ahmad FJ, Baas PW (1993) Microtubule nucleation and release from the neuronal centrosome. The Journal of cell biology 122: 349–359

