## Supplemental figures for "Centrosome Loss Alters Axonal and Muscle Development"

**Centrosome Loss in Embryonic Development Disrupts Axonal Pathfinding and Muscle Integrity**

### MATERIALS AND METHODS

#### Table S1

| Reagent type | Designation | Source | Additional information |
|---|---|---|---|
| Genetic reagent (*D. melanogaster*) | *Sas-4$^{s2214}$* | J. Raff | (Basto *et al*, 2006) |
| Genetic reagent (*D. melanogaster*) | *Sas-6$^{\Delta b}$* | J. Raff | (Zsofia Novak and Jordan Raff, unpublished) |
| Genetic reagent (*D. melanogaster*) | *UAS-Sas-4RNAi* | BDSC | BL35049 |
| Genetic reagent (*D. melanogaster*) | *UAS-Sas-4RNAi (2)* | VDRC | 106051 |
| Genetic reagent (*D. melanogaster*) | *UAS-AssRNAi* | VDRC | 47097 |
| Genetic reagent (*D. melanogaster*) | *pUb-Sas-4GFP* | J. Raff | (Cottee *et al*, 2013) |
| Genetic reagent (*D. melanogaster*) | *Df(3R) BSC221* | BDSC | BL9698 |
| Genetic reagent (*D. melanogaster*) | *Df(3R) BSC794* | BDSC | BL27366 |
| Genetic reagent (*D. melanogaster*) | *elav-Gal4* | BDSC | BL5146 |
| Genetic reagent (*D. melanogaster*) | *RN2-GAL4 UAS-CD8GFP* | BDSC | BL7472, BL7475 |
| Genetic reagent (*D. melanogaster*) | *Mef2-GAL4* | B. Estrada | (Pérez-Moreno *et al*, 2014) |
| Genetic reagent (*D. melanogaster*) | *Twi-GAL4* | B. Estrada | (Pérez-Moreno *et al.*, 2014) |
| Genetic reagent (*D. melanogaster*) | *P0163-GAL4* | C. Klambt | (Hummel *et al*, 2000) |
| Genetic reagent (*D. melanogaster*) | *ppkGAL4 UAS-CD8GFP* | S. Rumpf | (Herzmann *et al*, 2017) |
| Genetic reagent (*D. melanogaster*) | *AslYFP* | C. Gonzalez | (Varmark *et al*, 2007) |

| | | | |
|---|---|---|---|
| Genetic reagent (*D. melanogaster*) | *Tre-red* | E. Martin-Blanco | (Chatterjee & Bohmann, 2012) |
| Antibody | mouse anti FasII | DSHB | 1:5 |
| Antibody | mouse anti Futsch | DSHB | 1:10 |
| Antibody | rabbit anti CP309 (Plp) | Y. Zheng | 1:1000 (Kawaguchi & Zheng, 2004) |
| Antibody | goat, rabbit anti GFP | Roche / Jackson | 1:300 |
| Antibody | rabbit anti RFP | Abcam | 1:300 |
| Antibody | rabbit anti dsRed | Takara Bio | 1:600 |
| Antibody | chicken, rabbit, mouse anti-βgal | Cappel/ Promega/ Abcam | 1:300 |
| Antibody | mouse anti-acetylated tubulin | Sigma | 1:100 |
| Antibody | mouse anti-gamma tubulin | Sigma | 1:100 |
| Antibody | Rat anti-myosin | Abcam | 1:500 |
| Antibody | mouse biotinylated anti-IgG Secondary antibody | Vector Laboratories | 1:300 |
| Antibody | Cy2-conjugated secondary antibody | Jackson Labs | 1:500 |
| Antibody | Cy3-conjugated secondary antibody | Jackson Labs | 1:500 |
| Antibody | Cy5-conjugated secondary antibodies | Jackson Labs | 1:500 |
| Antibody | Alexa 488 conjugated | Thermo Fischer Scientific | 1:500 |

| | | | |
|---|---|---|---|
| | secondary antibodies | | |
| Antibody | Alexa 647 conjugated secondary antibodies | Thermo Fischer Scientific | 1:500 |
| Antibody | Alexa 555 conjugated secondary antibodies | Thermo Fischer Scientific | 1:500 |
| AB solution | Vectastain ABC kit | Vector Laboratories | 1:200 |

#### *Drosophila melanogaster* strains and genetics

Fly stocks described in Table S1 were maintained in plastic vials at 18°C or at 25°C on *Drosophila* culture medium (yeast 36%, glucose 36%, agar 5%, flour 22.5% and nipagin 1%). For microscopy, hatching and eclosion rates, and larval counting experiments, flies were placed in egg-laying cages on fruit juice plates (40% apple juice, 2% sucrose, and 1.8% agar) with a drop of yeast paste at 25°C.

Chromosomes were balanced over LacZ or GFP-labelled balancer chromosomes to differentiate homozygous mutant from heterozygous individuals. RNAi experiments were carried out either with *elav-GAL4, P0163-GAL4, Mef2-GAL4, TwiGAL4;Mef2GAL4* or *RN2-GAL4* at 25°C or 29°C as described. Rescues were performed at 25°C.

The control-RNAi line used in all the RNAi downregulation experiments is an RNAi against Argininosuccinate synthase (Ass-RNAi) a gene exclusively expressed in the Malpighian tubules (Dow & Romero, 2010). Other control lines are $w^{118}$, GAL4 only lines or heterozygous mutant embryos as described in the corresponding figure legends.

*Sas-6*$^{\Delta a}$ and *Sas-6*$^{\Delta b}$ mutants were generated by CRISPR-Cas9 in Jordan Raff's laboratory, Oxford University, UK. They are both deletion alleles induced by NHEJ in the *Sas-6* coding region (Zsofia Novak and Jordan Raff, unpublished). In order to avoid other genetic interferences we used the transallelic combination *Sas-6*$^{\Delta a}$/*Sas-6*$^{\Delta b}$ in this study.

#### Immunohistochemistry, image acquisition, and processing

All stage embryos, collected on agar plates overnight (o/n), were dechorionated with bleach for 2 min and fixed for 20 min (or 10 min for MT staining) in 4% formaldehyde, PBS (0.1 M NaCl, 10 mM phosphate buffer, pH 7.4) / Heptane 1:1). Washes were done with PBT (PBS, 0.1% Tween). Primary antibody incubation was performed in fresh PBT-BSA O/N at 4°C.

Secondary antibody incubations were done in fresh PBT-BSA at room temperature (RT) in the dark for 2h.

For DAB histochemistry (used to recognize anti-FasII and anti-Futsch antibody) after incubation with secondary antibody (anti-mouse IgG biotinylated antibody) embryos were treated with AB solution for 30 min at R/T (Avidin-Biotinylated Horseradish Peroxidase H from Vectastain-ABC KIT of Vector Laboratories 1:200 in PBT). Embryos were incubated with the DAB solution (DAB 0,12% Nickel Sulphate-Cobalt chloride, 0,3 %$H_2O_2$) until black colour was achieved, usually 3/5 min.

Bright field photographs were taken Nikon ECLIPSE Ci and LEICA DMLB fluorescence microscopes with a 20X or 40X objective. Fiji (Imagej 2.1 1.53c) (Schindelin J, 2012) was used for Z-projections measurements and adjustments. Photoshop v25 was used to assemble figures. Fluorescence confocal images of fixed embryos were obtained with Leica TCS-SPE system using 20X, 40X and 63X (1.40-0.60 oil) objectives (Leica). Fiji (Imagej 2.1 1.53c) was used for Z-projections, measurements and adjustments. The images shown are, otherwise stated in the text, Max-intensity projections of Z-stack sections.

**Quantitative Analysis**

Total number of embryos, segments and neurons quantified (n) are provided in the figure legends. The number of centrioles per aCC+pCC and RP2 neurons in embryos was counted in fixed and stained samples of stage 12-14 embryos. Axonal and muscle mutant phenotypes were scored in homozygous embryos derived from stocks carrying balancer chromosomes marked with either GFP or lacZ. Phenotypic scoring was conducted in a blind manner for both homozygous and heterozygous embryos. The genotype of each embryo, specifically the presence or absence of the balancer chromosome, was determined only after phenotypic evaluation was completed.

To quantify total centrioles, confocal images were acquired using a 63x objective. After image acquisition, the data were processed using the ImageJ software, and the centrioles, visualized as distinct punctate structures, were manually quantified. To do so, using ImageJ, the image channels were separated (in this study, two channels were analyzed, one containing the Z-stacks of the PNS and another containing the Z-stacks of the centrioles) to generate images displaying only the centrioles and the background was subtracted using a rolling ball radius of 10 pixels. For quantification, a specific region of the PNS was consistently selected, delineated by a rectangle measuring 618x260 pixels in six different segments per embryo, and all centrioles within this area were counted. To ensure accuracy and avoid duplication, each centrosome was individually marked with a colour distinct from that of the original image. This procedure was repeated for all embryos, and the resulting data were subjected to statistical analysis. Axonal wiggliness axonal wiggliness via the ratio of Path Length in 2D (PL2D) normalized to Euclidean Distance in 2D (PL2D/ED2D) was quantified using the plugin "SNT" from Fiji/ImageJ (Arshadi *et al*, 2021).

**Statistical analysis**

Measurements were imported and treated in GraphPad Prism version 10.4, where graphs were generated, and all statistical analysis were performed.

The data were tested for normal distribution using the Shapiro-Wilk test. When the data followed a normal distribution and only one experimental condition was compared with a control, an unpaired Student's t-test with Welch's correction was performed. When the data did not follow a normal distribution, a Mann-Whittney U-test was performed. Differences in the distribution of phenotypic categories between groups were assessed using a Pearson's chi-square test. Statistical significance was determined at $p < 0.05$. Error bars in graphs and ± in text denote Standard deviation (SD). Statistical significance was defined as $p<0.05$. In graphics; *p<0.05, **p<0.01, ***p< 0.001, **** p< 0.0001; ns is p>0.05.

**Supplementary figure legends**

**Supplementary Fig. 1 - *Sas-4* loss of function induces axonal misrouting defects**

(A-E) Details of four segments from stage 16 embryos stained with anti-FasII antibody and analysed by light microscopy, using colorimetric methods. This allows us to quantify the whole embryonic phenotypes, from the two embryonic laterals; scale bar 10 μm. (F-G) Details of four segments from stage 16 embryos stained with anti-Futsch antibody and analysed by light microscopy, using colorimetric methods; scale bar 10 μm (K) Quantification of ISN phenotypes. (L) Quantification of sensory axon phenotypes.

(M-O) Analysis of motor axon guidance phenotypes by Sas-4 downregulation using an alternative RNAi line (Sas-4RNAi 2). (M) RN2GAL4 control; (N,O) Sas-4 downregulation. Embryos were stained with anti-FasII antibody, and images are confocal projections; scale bar is 10 μm.

Anterior is left and dorsal is up in all images.

**Supplementary Fig. 2 - *Sas-4* mutant conditions induce muscle developmental defects**

(A, B) Confocal projections of three segments from stage 16 embryos of control and *Sas-4* mutant embryos stained with anti-myosin (green and A, B) and anti-Futsch (magenta and A', B') antibodies. (A'', B'') Composite images with the two channels; scale bar 10 µm. (D, E) Confocal projections of three segments from stage 16 embryos of control (*Twi,Mef2Gal4*) and *Twi,Mef2Gal4xUASSas-4RNAi* embryos stained with anti-myosin (green and D, E) and anti-Futsch (magenta and D', E') antibodies. (D'', E'') Composite images with the two channels; scale bar 10 µm. (F) Quantification of the ISN axon phenotypes in the embryos expressing *Sas-4RNAi* in the muscle.

**Supplementary Fig. 3 – Asl and Plp colocalize in neurons**

(A, A') Confocal projection of stage 12 FasII positive neurons (blue) at the VNC from an embryo expressing Asl-YFP (green) and stained using an antibody against Plp (CP309, magenta). Scale bar is 5 µm. Three neurons are depicted, and arrows point at the centrosomes. (B) Quantification of the number of Asl, Plp and Asl and Plp positive centrosomes in neurons at the VNC (84.4% colocalization, n=60 neurons).

**Supplementary Fig. 4 – JNK signaling is not activated in *Sas-4* VNC neurons**

(A) Confocal projection of stage 13 FasII positive neurons (green) at the VNC from an embryo expressing TREred (magenta), a reporter of JNK activity (Chatterjee & Bohmann, 2012). No JNK activity was detected in neurons in control and mutant conditions, despite being detected strongly at the leading-edge cells of the dorsal epidermis (not shown). Asterisk mark wiggly axon, scale bars are 5 µm.

**Supplementary references**


Arshadi C, Günther U, Eddison M, Harrington KIS, Ferreira TA (2021) SNT: a unifying toolbox for quantification of neuronal anatomy. *Nature Methods* 18: 374-377

Basto R, Lau J, Vinogradova T, Gardiol A, Woods CG, Khodjakov A, Raff JW (2006) Flies without centrioles. *Cell* 125: 1375-1386

Chatterjee N, Bohmann D (2012) A versatile PhiC31 based reporter system for measuring AP-1 and Nrf2 signaling in Drosophila and in tissue culture. *PLoS One* 7: e34063

Cottee MA, Muschalik N, Wong YL, Johnson CM, Johnson S, Andreeva A, Oegema K, Lea SM, Raff JW, van Breugel M (2013) Crystal structures of the CPAP/STIL complex reveal its role in centriole assembly and human microcephaly. *eLife* 2: e01071
Dow JAT, Romero MF (2010) Drosophila provides rapid modeling of renal development, function, and disease. *Am J Physiol-renal* 299: F1237-F1244
Herzmann S, Krumkamp R, Rode S, Kintrup C, Rumpf S (2017) PAR-1 promotes microtubule breakdown during dendrite pruning in Drosophila. *The EMBO Journal* 36: 1981-1991
Hummel T, Krukkert K, Roos J, Davis G, Klämbt C (2000) Drosophila Futsch/22C10 Is a MAP1B-like Protein Required for Dendritic and Axonal Development. *Neuron* 26: 357-370
Kawaguchi S-i, Zheng Y (2004) Characterization of a Drosophila centrosome protein CP309 that shares homology with Kendrin and CG-NAP. *Mol Biol Cell* 15: 37-45
Pérez-Moreno JJ, Bischoff M, Martín-Bermudo MD, Estrada B (2014) The conserved transmembrane proteoglycan Perdido/Kon-tiki is essential for myofibrillogenesis and sarcomeric structure in Drosophila. *Journal of Cell Science* 127: 3162-3173
Varmark H, Llamazares S, Rebollo E, Lange B, Reina J, Schwarz H, Gonzalez C (2007) Asterless is a centriolar protein required for centrosome function and embryo development in Drosophila. *Curr Biol* 17: 1735-1745
